# GPNMB enables lysosomal adaptation required for myelin clearance and nerve regeneration after peripheral nerve injury

**DOI:** 10.64898/2026.10.12.758439

**Authors:** Najwa Ouali Alami, Moh Panesar, Michael Obrecht, Thierry Doll, Ivan Osinni, Gilles Sansig, Zacharias Thiel, Isabelle Claerr, Daniel Greenwood, Claudia Patino, Laurent Klein, Daniela Condeescu-Ivan, Elisa Giorgetti, Christian Lambert, Georgios Kalamakis, Valerie Van Steenbergen, Lukas Hils, Carole Manneville, Matthias Müller, Mark Nash, Marie Labouesse, Derya Shimshek, Nicolau Beckmann, Oleksandr Dergai, Michael Bidinosti

## Abstract

Recovery from peripheral nerve injury involves coordinated axonal and immune repair programs that promote tissue regeneration. However, the mechanisms regulating immune-mediated repair remain incompletely understood. Here, we demonstrate that glycoprotein non-metastatic melanoma protein B (GPNMB) is a key regulator of repair processes following peripheral nerve injury. In a mouse sciatic nerve crush model, loss of *Gpnmb* impaired functional recovery after injury. MRI and histological analyses revealed persistent myelin debris accumulation, defective debris clearance, and impaired macrophage lysosomal function within injured nerves. Mechanistic studies showed that GPNMB-deficient human iPSC-derived macrophages exhibited impaired lysosomal adaptation, reduced lysosomal fitness, and defective processing of myelin-associated cargo, resulting in inefficient degradation of injury-associated material. These defects were accompanied by impaired acquisition of repair-associated cellular programs. Together, our findings identify GPNMB as a critical regulator of lysosome-dependent repair programs during nerve regeneration and demonstrate that efficient GPNMB-dependent myelin processing is required for functional recovery following peripheral nerve injury.

**Graphical abstract:** **GPNMB enables efficient nerve regeneration after injury**

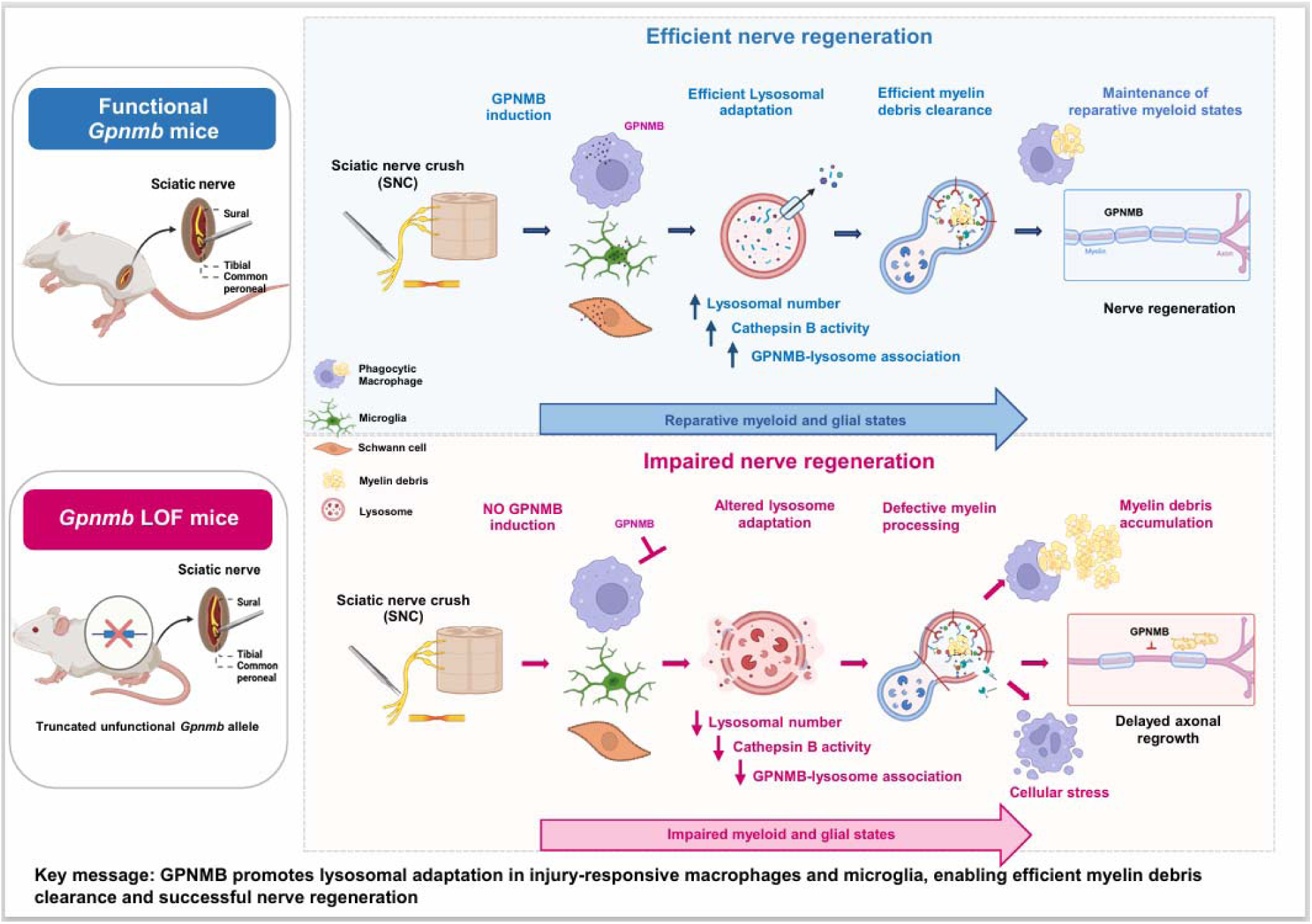

**GPNMB promotes lysosomal adaptation in injury-responsive macrophages and microglia, enabling efficient myelin debris clearance and successful nerve regeneration.**

GPNMB coordinates injury-responsive myeloid and glial remodeling following peripheral nerve injury. After sciatic nerve crush, GPNMB is induced in macrophages, Schwann cells, and spinal cord microglia, where it promotes lysosomal adaptation and efficient myelin debris processing. By supporting reparative cellular states and sustained debris clearance, GPNMB enables axonal regeneration and functional recovery.

## Introduction

Damage and degeneration of lower motor neuron axons are central features of multiple neurodegenerative diseases, including amyotrophic lateral sclerosis (ALS), diabetic neuropathy, and Charcot–Marie–Tooth disease (CMT), as well as acute peripheral nerve injury^1–3^. In these conditions, disruption of axonal integrity leads to loss of neuromuscular connectivity and functional decline. Axonal degeneration is driven by convergent molecular mechanisms, including activation of the SARM1 pathway, mitochondrial dysfunction, impaired axonal transport, and cytoskeletal breakdown^4–6^.

Peripheral nerve repair is critically dependent on coordinated immune and glial responses. Following injury, Wallerian degeneration triggers activation of resident macrophages and Schwann cells and recruitment of circulating immune cells to mediate myelin and cellular debris clearance^7,8^. This initial inflammatory phase is followed by a transition toward pro-repair states, characterized by Schwann cell reprogramming and the emergence of reparative macrophages that promote axon growth, tissue remodeling, and restoration of homeostasis^9^. This phenotypic shift from pro-inflammatory to pro-regenerative states is a key determinant of regeneration success. Notably, the response to peripheral nerve injury is not restricted to the injured nerve but extends to the central nervous system (CNS). Sciatic nerve injury induces retrograde signaling that activates spinal cord circuits and promotes microglial proliferation and phenotypic changes^10–14^. These microglial responses include the release of soluble factors that can support regeneration but also contribute to central sensitization and neuropathic pain^15,16^. Thus, peripheral nerve injury engages a coordinated PNS–CNS immune response.

GPNMB is a transmembrane glycoprotein expressed in several cell types including macrophages, microglia, and other immune cells and has been shown to have key functions in immune modulation and neuroprotection, in part through cellular regulation of phagocytosis, autophagy and lysosomal function^17–20^. GPNMB can also be proteolytically cleaved to generate a soluble fragment capable of mediating autocrine and paracrine signaling^21^. In macrophages and microglia, GPNMB expression is low under homeostatic conditions but is robustly induced in inflammatory and neurodegenerative settings, including Alzheimer’s disease, ALS, and Parkinson’s disease^22–24^. In the CNS, GPNMB is highly expressed in disease-associated microglia (DAM), a state characterized by enhanced phagocytic activity, increased lysosomal function, and reduced inflammatory signaling^19,25^. However, whether GPNMB actively drives the acquisition and function of these injury-or disease-associated myeloid states, or simply reflects the underlying cellular state, remains unclear. GPNMB expression is also induced following peripheral nerve injury. Single-cell transcriptomic analyses reveal that *Gpnmb*⁺ macrophages are absent from uninjured peripheral nerves but expand significantly during Wallerian degeneration, marking mature injury-associated macrophage states^26^. GPNMB expression is also detected in Schwann cells and increases in distal nerve segments after injury, with peak expression coinciding with debris clearance and regeneration^27,28^. Moreover, GPNMB is enriched in macrophage subsets associated with lipid handling and tissue repair and is functionally linked to pro-regenerative responses^29,30^.

Together, these observations suggest that GPNMB contributes to recovery following peripheral nerve injury; however, direct causal evidence for this role is lacking. Here, we addressed this question using a sciatic nerve crush model in adult mice, combined with in vivo functional and imaging analyses and in vitro mechanistic studies in human cell systems. We identify GPNMB as a central regulator of injury-induced myeloid cell responses that promotes lysosomal integrity, myelin debris clearance, and axon regeneration.

## Results

### *Gpnmb* deficiency delays neuromuscular recovery after sciatic nerve crush

We first examined whether GPNMB expression is induced following peripheral nerve injury. To do so, we reanalyzed published mouse transcriptomic datasets^31,32^ and generated a de novo longitudinal rat sciatic nerve crush RNA-seq dataset (**Fig. S1A-S1C**). Across all datasets, *Gpnmb* expression was robustly upregulated following injury and temporally coincided with the period of active axonal regeneration, as indicated by increased *Stmn2* expression (**Fig. S1D-E**). Immunofluorescence analyses confirmed strong GPNMB expression during this phase in both IBA1⁺ macrophages (**Fig. S1F)** and S100β⁺ Schwann cells within injured nerves compared with sham-operated controls (**Fig. S1G)**. Strikingly, *Gpnmb* induction was largely absent in Wallerian degeneration slow (Wld^S^) mice^31^, which, following injury, exhibit profoundly delayed axonal degeneration and attenuated injury-associated molecular and cellular responses (**Fig. S1B**). These findings indicate that GPNMB induction depends on functional axon degeneration and suggest that GPNMB may act as a molecular link between Wallerian degeneration and successful nerve regeneration.

Because peripheral nerve injury also elicits microglial responses in the central nervous system^11,33,34^, we next examined GPNMB expression in the spinal cord. GPNMB expression was markedly increased in IBA1-positive microglia between 3 and 14 days post-crush (dpc), coinciding with microglial proliferation and morphological remodeling in both the dorsal and ventral spinal cord regions (**Fig. S2A-I**). GPNMB-positive microglia were frequently observed in close proximity to motor neuron cell bodies^35^ during the regenerative phase following injury (**Fig. S2J-K**). Thus, GPNMB is robustly induced in injury-responsive myeloid populations in both the peripheral nerve and spinal cord following sciatic nerve injury. This supports a key role for GPNMB in the cellular repair processes that enable nerve regeneration.

We therefore investigated whether GPNMB plays a causal role in recovery following peripheral nerve injury using our established sciatic nerve crush model^36^. DBA/2J mice carrying a spontaneous loss-of-function mutation in *Gpnmb* (*Gpnmb*^R150X^; hereafter referred to as *Gpnmb*-null) were compared with isogenic controls expressing intact *Gpnmb* (*Gpnmb*-WT). Animals were subjected to sciatic nerve crush and monitored longitudinally for functional recovery (**Fig. 1A**).

**Figure 1.**
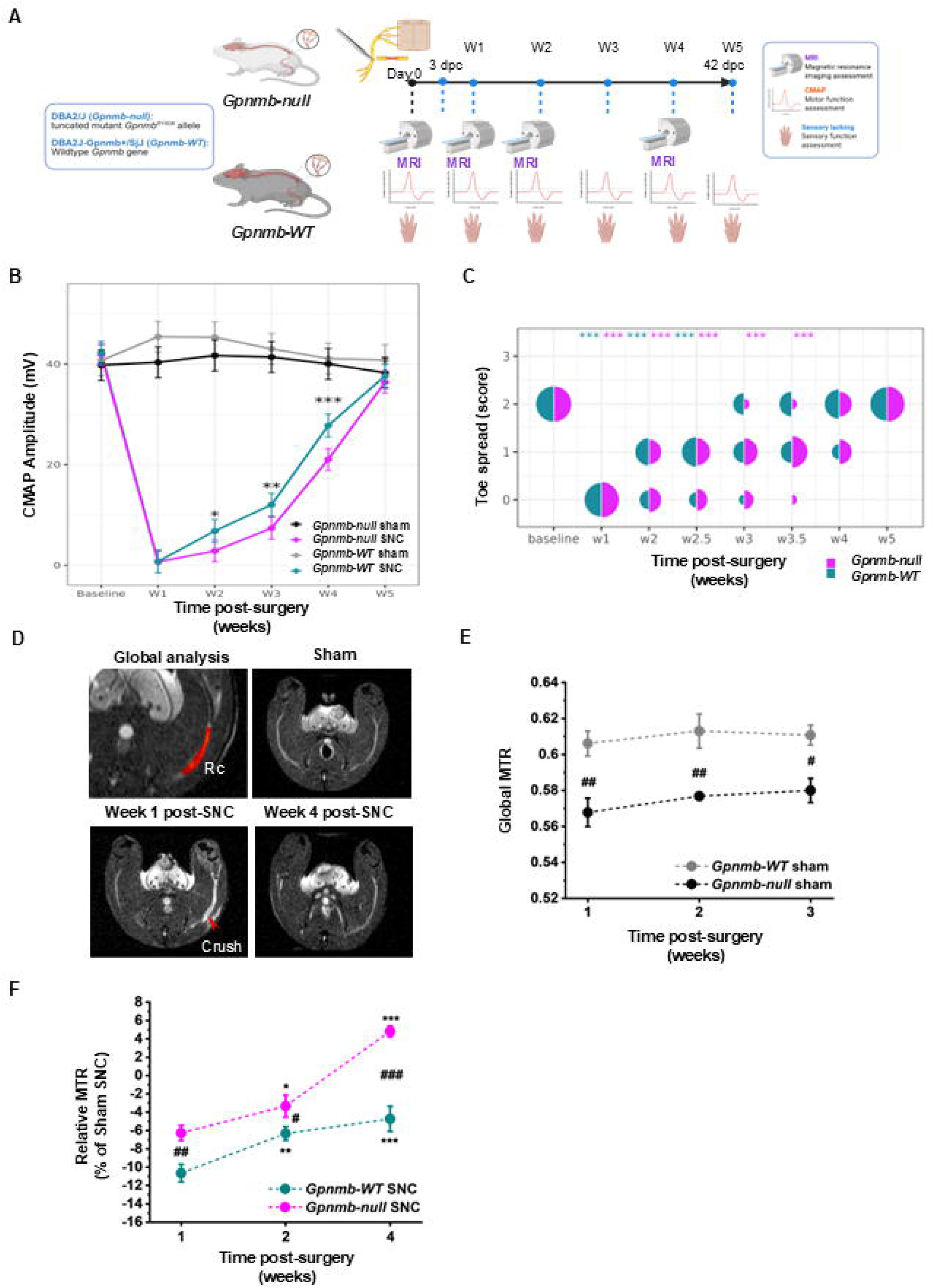
***Gpnmb* deficiency delays functional recovery and alters peripheral nerve remodeling after sciatic nerve injury** **(A)** Experimental design. Adult *Gpnmb*-null (DBA2/J) and *Gpnmb*-wild-type (DBA2J-Gpnmb+/SjJ) mice underwent unilateral sciatic nerve crush (SNC). Functional recovery was monitored longitudinally by compound muscle action potential (CMAP) recordings and toe spread (TS) assessment. Nerve remodeling was evaluated by magnetic resonance imaging (MRI) at the indicated time points following injury (CMAP and TS in week 0, 1, 2, 3, 4 and 5; MRI in week 0, 1, 2 and 4). **(B)** Longitudinal assessment of CMAP amplitude following sciatic nerve crush injury. CMAP amplitudes were markedly reduced in both genotypes at week 1 (W1) post injury (wpi) and gradually recovered over subsequent weeks. Statistical significance was determined using a linear mixed-effects model including strain, timepoint, surgery, and their interactions, followed by estimated marginal means (EMMs) and Holm-corrected pairwise comparisons between genotypes within the sciatic nerve crush group at each timepoint. **(C)** Toe spread recovery following sciatic nerve crush in *Gpnmb*-null and *Gpnmb*-WT mice. Split-circle bubble plots depict the distribution of ipsilateral toe spread scores following sciatic nerve crush (SNC) in *Gpnmb*-null (magenta, left semicircle) and *Gpnmb-*WT (teal, right semicircle) mice across time. Toe spread was scored as 0, absent; 1, partial; and 2, full toe spreading. Circle area is proportional to the number of animals at each score. Significance markers above each semicircle indicate time points at which ipsilateral toe spread scores differed significantly from the corresponding contralateral (uninjured) limb within each genotype (Wilcoxon rank-sum (Mann-Whitney) test; *p* < 0.05, p < 0.01, *p* < 0.001). See Materials and Methods for additional statistical details. **(D)** Representative axial magnetization transfer (MT) MRI images of sham-operated nerves and injured nerves at weeks 1 and 4 following SNC. The crush site is indicated by the red arrow. **(E)** Global magnetization transfer ratio (MTR) measured across the entire sciatic nerve segment in sham-operated *Gpnmb*-null and *Gpnmb*-WT mice at time points corresponding to weeks 1, 2, and 4 after surgery. **(F)** Relative change in global MTR (ΔMTR) of the sciatic nerve segment following SNC, expressed as the percentage difference from genotype-matched sham controls at the corresponding time points. Negative values indicate a reduction in MTR relative to sham-operated animals. Data are presented as mean ± SEM. Statistical analysis was performed using a random-effects ANOVA followed by Fisher’s post hoc test. *P* values are denoted as *p < 0.05, **p < 0.01, and ***p < 0.001 versus week 1 within the same genotype; #p < 0.05, ##p < 0.01, and ###p < 0.001 between genotypes at the indicated time points. TS: Statistical significance was assessed using repeated-measures ordinal regression with genotype, treatment, time, and their interactions included in the model, followed by multiple-comparison testing.

Compound muscle action potential (CMAP) recordings, which provide a measure of muscle reinnervation, revealed significantly delayed neuromuscular recovery in *Gpnmb*-null mice following injury. In particular, CMAP amplitudes remained significantly reduced at 3 and 4 weeks post-injury, relative to *Gpnmb*-WT controls (**Fig. 1B**). Consistent with these findings, assessment of toe-spread reflex recovery showed a similar trend toward delayed functional recovery in *Gpnmb*-null mice. Although both genotypes exhibited complete loss of toe-spread responses one week after injury, recovery was slower in *Gpnmb*-null animals from approximately 3 weeks post-injury onward (**Fig. 1C**). At these later timepoints, a greater proportion of *Gpnmb*-WT mice regained full toe-spread function, whereas recovery remained incomplete in several *Gpnmb*-null mice. Thus, loss of GPNMB impairs functional recovery following peripheral nerve injury.

To assess injury-induced changes in nerve tissue remodeling, we also longitudinally profiled peripheral nerves using magnetic resonance imaging (MRI). Magnetization transfer ratio (MTR) imaging, a quantitative measure sensitive to tissue macromolecular composition and widely used as a surrogate marker of myelin integrity and content in peripheral nerves^36,37^, was acquired at 1, 2, and 4 weeks following sciatic nerve crush (**Fig. 1A**). Global MTR analysis of the injured nerve segment (**Fig. 1D**) revealed lower MTR values in sham-operated *Gpnmb*-null mice compared with isogenic controls, indicating differences in nerve tissue composition under uninjured conditions (**Fig. 1E**). Following sciatic nerve crush, both genotypes exhibited a sharp reduction in MTR at week 1, consistent with injury-induced demyelination (**Giorgetti, E., et al., 2019**), However, the reduction in MTR was significantly greater in *Gpnmb*-WT mice (∼10.6%) than in *Gpnmb*-null mice (∼6.3%) relative to their respective sham controls (**Fig. 1F**). Surprisingly, from 2 to 4 weeks post-injury, relative MTR values increased significantly more in *Gpnmb*-null nerves than in control nerves (**Fig. 1F; Fig. S3A-B**). In contrast, spinal cord MTR values were similar between genotypes at all time points examined (**Fig. S3C-E**).

T2-weighted MRI, which reflects injury-associated tissue changes including edema and inflammation, revealed a transient increase in signal intensity of the sciatic nerve during the acute post-injury phase, with values returning toward baseline by 4 weeks in both genotypes (**Fig. S3F**). Together, these findings demonstrate that loss of GPNMB is associated with delayed functional recovery and altered tissue remodeling following peripheral nerve injury. Notably, genotype-dependent MRI abnormalities were restricted to the injured sciatic nerve, in comparison to spinal cord, suggesting that local changes within the regenerating peripheral nerve contribute to the impaired recovery phenotype observed in *Gpnmb*-null mice.

### Loss of *Gpnmb* causes myelin debris accumulation and altered autophagy– lysosomal processing

Because delayed neuromuscular recovery and alterations in longitudinal MRI imaging suggested impaired tissue repair in *Gpnmb*-deficient nerves, we next asked whether loss of GPNMB compromises myelin debris clearance after injury. We quantified degraded myelin basic protein (dMBP) in distal sciatic nerve segments at 3 and 42 days post-crush (dpc) (**Fig. 2A-B**). While both genotypes exhibited dMBP accumulation shortly after injury, dMBP levels were elevated in *Gpnmb*-deficient nerves at 3 dpc. This difference was even more pronounced at 42 dpc, when dMBP had been largely cleared from *Gpnmb*-WT nerves but remained persistently elevated in *Gpnmb*-deficient animals (**Fig. 2B-C**), indicating defective clearance of injury-derived myelin debris in the absence of GPNMB.

**Figure 2.**
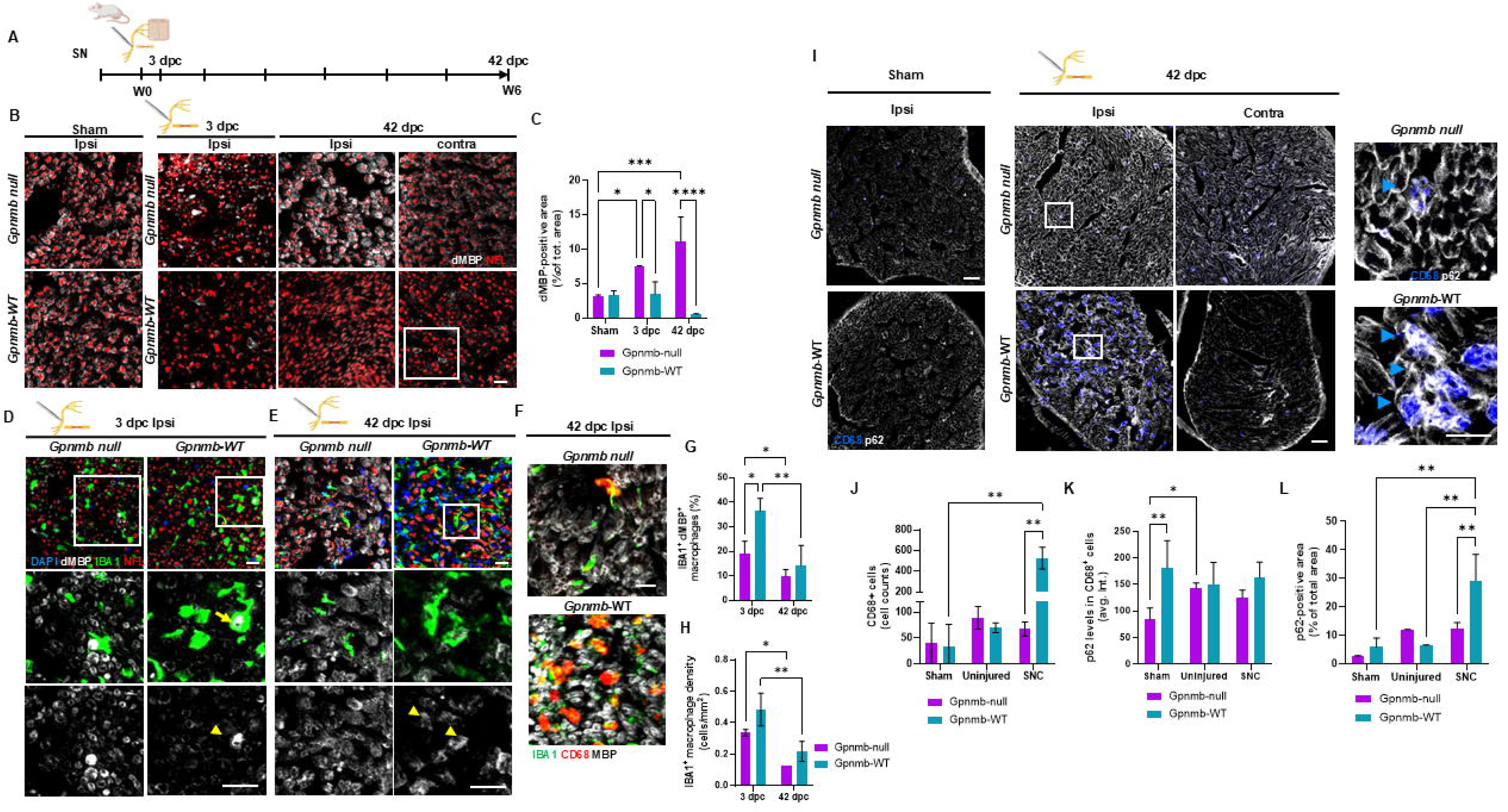
***Gpnmb* loss impairs myelin clearance, injury-associated macrophage activation, and tissue remodeling after nerve injury** (A) Experimental overview. Sciatic nerve tissues were collected from *Gpnmb*-null and wild-type mice at 3 and 42 days post-crush (dpc) for histological analyses. (B) Representative confocal images of transverse sciatic nerve sections from sham-operated mice and injured nerves collected at 3 and 42 days post-crush (dpc), stained for degraded MBP (dMBP, gray) and neurofilament light chain (NFL, red). Increased accumulation of dMBP-positive material was observed in *Gpnmb*-null nerves compared with wild-type controls following injury. Scale bar, 50 μm. (C) Quantification of dMBP-positive area in sciatic nerves from *Gpnmb*-WT and *Gpnmb*-null mice at 3 and 42 dpc. Data are presented as mean ± SD. Statistical analysis was performed using a mixed-effects model (REML) including genotype, time, and genotype × time interaction terms, followed by Šidák’s multiple-comparisons test. (D) Representative confocal images of transverse sciatic nerve sections from *Gpnmb*-null and wild-type at 3 dpc stained for degraded MBP (dMBP, gray), IBA1 (green), NFL (red), and DAPI (blue). Insets show higher-magnification views of myelin-associated IBA1-positive macrophages (arrows). Scale bars, 50 μm (overview images) and 15 μm (magnified views). (E) Representative confocal images of transverse sciatic nerve sections from *Gpnmb*-null and wild-type at 42 dpc stained for degraded MBP (dMBP, gray), IBA1 (green), NFL (red), and DAPI (blue). Insets show higher-magnification views of myelin-associated IBA1-positive macrophages (arrows). Scale bars, 50 μm (overview images) and 15 μm (magnified views). (F) Representative confocal images of transverse sciatic nerve sections from *Gpnmb*-null and wild-type mice collected at 42 dpc and stained for IBA1 (green), CD68 (red), and MBP (gray). Scale bar, 40 μm. (G) Quantification of IBA1⁺ macrophages containing dMBP-positive myelin debris, expressed as a percentage of total IBA1⁺ macrophages in the distal sciatic nerve at 3 and 42 dpc. Data are presented as mean ± SD. Statistical analysis was performed using a mixed-effects model (REML) including genotype, time, and genotype × time interaction terms, followed by Fisher’s LSD test (uncorrected). (H) Quantification of IBA1⁺ macrophage density in the distal sciatic nerve at 3 and 42 dpc. Data are presented as mean ± SD. Statistical analysis was performed using a mixed-effects model (REML) including genotype, time, and genotype × time interaction terms, followed by Fisher’s LSD test (uncorrected). (I) Representative confocal images of transverse sciatic nerve sections from wild-type and *Gpnmb*-null mice at 42 dpc stained for CD68 (blue) and p62 (gray). Boxed regions are shown at higher magnification in the corresponding right panels. Arrowheads indicate representative p62-positive structures associated with CD68⁺ macrophages. Scale bars, 50 μm (main panels) and 10 μm (high-magnification images). (J) Quantification of CD68-positive cells within the endoneurium of contralateral (uninjured) and ipsilateral (injured, SNC) sciatic nerves from *Gpnmb*-null and wild-type mice at 42 dpc. Data are presented as mean ± SD. Statistical analysis was performed using a mixed-effects model (REML) with genotype, injury status, and genotype × injury status interaction terms, followed by Šidák’s multiple-comparisons test. (K) Quantification of p62 mean intensity associated with CD68⁺ macrophages in transverse sciatic nerve sections from wild-type and *Gpnmb*-null mice at 42 dpc. (L) Quantification of p62-positive area normalized to total nerve area in injured *Gpnmb*-null and wild-type sciatic nerves at 42 dpc. Data are presented as mean ± SD. Statistical analysis was performed using two-way ANOVA followed by Tukey’s multiple-comparisons test. Data are presented as mean ± SD. Statistical analyses were performed using one-way ANOVA, two-way ANOVA, or mixed-effects models (REML) with genotype, time, and genotype × time interaction terms. For analyses in panels C, G, and H, three wild-type and two *Gpnmb*-null mice were analyzed. Measurements from multiple images were averaged to generate a single value per animal, which was then used as the unit of statistical analysis. Mixed-effects models were followed by Šidák’s multiple-comparisons test (C) or uncorrected Fisher’s LSD test (G, H). *p* < 0.05, p < 0.01, *p* < 0.001, p < 0.0001.

In contrast to dMBP, *Gpnmb*-null mice exhibited reduced total MBP staining in uninjured nerves and 3 dpc compared with WT controls. However, by 42 dpc, MBP staining had increased relative to 3 dpc and was comparable to that observed in WT nerves (**Fig. S4A-B**). Because the MBP antibody recognizes epitopes present in both intact myelin and myelin debris^38,39^, the increased MBP staining in *Gpnmb*-deficient nerves is consistent with the elevated MRI signal and dMBP levels and may reflect newly formed myelin, persistent MBP-positive debris, or a combination of both.

To further assess axonal remodeling, we quantified neurofilament light chain (NFL), a key component of the axonal cytoskeleton. NFL-positive area was reduced at 3 dpc relative to sham nerves in both genotypes and showed a trend toward increased levels by 42 dpc (**Fig. S4C**). Although NFL-positive area remained lower than in sham nerves, WT mice tended to exhibit greater recovery than *Gpnmb*-deficient mice at the later time point. Together, these findings suggest altered myelin and axonal remodeling dynamics in the absence of GPNMB following peripheral nerve injury.

To determine whether myelin debris accumulation was associated with altered macrophage handling of myelin cargo, we examined the localization of dMBP relative to IBA1-positive macrophages. In control nerves, intracellular dMBP immunoreactivity was detected within a proportion of macrophages at 3 dpc, consistent with the uptake of degenerating myelin during Wallerian degeneration (**Fig. 2D, G**). In contrast, *Gpnmb*-deficient nerves contained fewer macrophages with intracellular dMBP inclusions. By 42 dpc, dMBP signal was largely resolved in control nerves but remained abundant in *Gpnmb*-deficient nerves, with limited evidence of macrophage-associated debris internalization (**Fig. 2E, G**).

The persistence of myelin debris was accompanied by marked alterations in the macrophage response. *Gpnmb*-deficient nerves displayed reduced IBA1-positive cell coverage and a reduction in macrophage size compared with control nerves (**Fig. 2E–F, H**). In addition, CD68 expression was reduced, consistent with diminished acquisition of a degradative, lysosome-rich macrophage state (**Fig. 2F**). Loss of GPNMB therefore impairs myelin debris clearance and macrophage remodeling after nerve injury, providing a potential explanation for the delayed functional recovery observed in *Gpnmb*-deficient mice.

Myelin accumulation in *Gpnmb*-deficient nerves suggested impaired cellular processing of injury-derived debris. Given the reported role of GPNMB in autophagy and lysosomal clearance pathways, we examined p62/SQSTM1, an autophagy adaptor that accumulates in response to autophagic and lysosomal stress^40,41^. At 42 dpc, injured nerves from *Gpnmb*-WT mice showed prominent p62 immunoreactivity within CD68⁺ macrophages. In *Gpnmb*-deficient nerves, CD68⁺ macrophages were markedly reduced in number and exhibited a smaller morphology compared with those in WT nerves, while retaining detectable p62 immunoreactivity (**Fig. 2I-J**).

Although p62 immunoreactivity was detected within CD68⁺ macrophages in both genotypes (**Fig. 2I-K**), *Gpnmb*-deficient injured nerves contained markedly fewer and morphologically smaller CD68⁺ macrophages than WT nerves. This reduction occurred despite the persistence of myelin debris in the injured nerve. Quantification further showed reduced overall p62-positive area in *Gpnmb*-null nerves (**Fig. 2I-L**), likely reflecting the decreased abundance of macrophages. Together, these findings indicate that GPNMB promotes the accumulation of enlarged injury-associated macrophages that support efficient myelin clearance during nerve repair.

To determine whether the effects of GPNMB deficiency extended beyond the injured nerve, we examined spinal cord microglia at 3 and 42 days post-crush (dpc). In control animals, sciatic nerve injury elicited a robust microglial response in the ventral horn, characterized by increased IBA1-positive coverage, elevated CD68 expression, and acquisition of an enlarged, hypertrophic morphology (**Fig. S5A-G**). CD68 induction was particularly prominent in microglia surrounding injured motor neurons, consistent with the emergence of lysosome-rich, injury-responsive microglial states (**Fig. S5F-G**). In contrast, *Gpnmb*-deficient mice exhibited an attenuated microglial response, characterized by significantly fewer IBA1⁺ cells (**Fig. S5B**), markedly reduced CD68 expression (**Fig. S5C**), and smaller microglial somata (**Fig. S5D-E**), consistent with impaired acquisition of an injury-associated activated phenotype. Notably, GPNMB colocalized with IBA1⁺ microglia, and although reduced, GPNMB immunoreactivity remained detectable in IBA1⁺ microglia at 42 days following injury (**Fig. S5H**). GPNMB therefore modulates the acquisition of injury-associated myeloid states in both the injured peripheral nerve and the spinal cord following sciatic nerve injury.

Collectively, our data suggest that GPNMB promotes efficient autophagy–lysosomal processing and the acquisition of injury-responsive myeloid states during nerve repair. In its absence, injury-derived myelin debris persists, p62 handling becomes altered, and both peripheral macrophages and spinal cord microglia fail to fully acquire the phagolysosomal phenotype associated with efficient myelin debris clearance and peripheral nerve repair.

### GPNMB expression positively correlates with lysosomal and lipid-processing programs in injury-associated macrophages

Given that *Gpnmb*-deficient nerves show impaired myelin debris clearance and reduced acquisition of CD68-positive macrophage states, we next asked whether GPNMB expression defines a distinct macrophage population specialized for tissue repair. To address this, we reanalyzed a published single-cell RNAseq dataset of injured sciatic nerve^26^. Across all cell populations, *Gpnmb* expression was dynamically regulated following injury and became increasingly enriched within macrophage populations during the regenerative phase (**Fig. 3A**). The highest proportion of *Gpnmb*-expressing cells was observed in injury-responsive macrophage clusters, suggesting that GPNMB is associated with specialized macrophage states that emerge during nerve repair.

**Figure 3.**
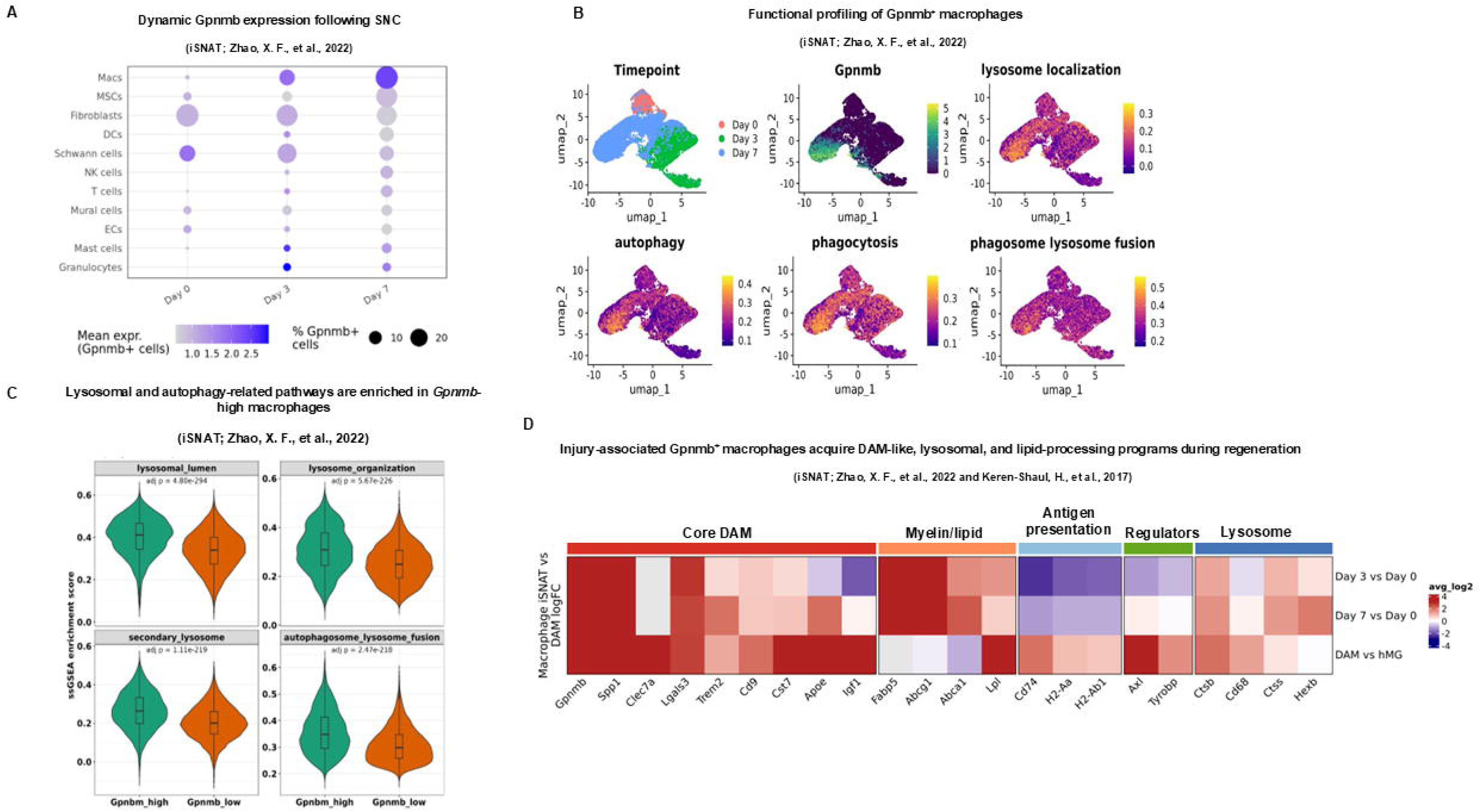
GPNMB marks injury-associated macrophages enriched for lysosomal, lipid-processing, and reparative gene programs. **(A)** Dynamic expression of Gpnmb across major cell populations following sciatic nerve crush. Dot plot showing the percentage of Gpnmb-expressing cells (dot size) and mean Gpnmb expression levels (color intensity) at Day 0, Day 3, and Day 7 post-injury. Macrophages exhibited the highest induction of Gpnmb expression following injury. **(B)** UMAP visualization of macrophages from the injured sciatic nerve atlas (iSNAT) dataset (Zhao et al., 2022), colored by time point after injury (top left), *Gpnmb* expression (top right), and single-cell Gene Set Enrichment Analysis (scGSEA) scores for autophagy (bottom left) and phagocytosis (bottom right) gene sets. **(C)** Violin plots showing ssGSEA enrichment scores for the four lysosome- and autophagy-related gene sets most significantly enriched in the *Gpnmb*-high macrophage cluster (cluster 1) relative to all other macrophage clusters (*Gpnmb*-low clusters). Gene sets are ranked by adjusted *p* value from a generalized linear model comparing *Gpnmb*-high and *Gpnmb*-low cells and include lysosomal lumen, lysosome organization, secondary lysosome, and autophagosome-lysosome fusion. Overlaid boxplots indicate the median and interquartile range. Adjusted *p* values (Benjamini-Hochberg correction) are shown above each panel. **(D)** Heatmap showing the expression of genes associated with antigen presentation, lysosomal function, transcriptional regulation, myelin/lipid processing, and core disease-associated migroglia (DAM)-related programs across macrophage subpopulations. *Gpnmb*⁺ macrophages at days 3 and 7 following injury displayed increased expression of lysosomal genes (*Cd68, Ctsb, Lyz2*), lipid-processing genes (*Apoe, Fabp5, Abca1*), and DAM-associated markers (*Trem2, Lpl, Spp1, Gpnmb*). The bottom row shows DAM vs homeostatic microglia expression changes for comparison. Color intensity represents normalized gene expression or single-cell GSEA enrichment scores as indicated. Dot size corresponds to the percentage of expressing cells.

To define the molecular programs associated with GPNMB expression, we compared *Gpnmb*-high and *Gpnmb*-low macrophage populations. Gene ontology analysis identified enrichment of lysosomal, autophagic, phagocytic, lipid-processing, and tissue-remodeling pathways in *Gpnmb*-expressing macrophages (**Fig. S6A**). Consistent with these findings, *Gpnmb*^+^ macrophages localized to regions enriched for lysosome-related, autophagic, phagocytic, and phagosome-lysosome fusion signatures (**Fig. 3B**). Single-sample Gene Set Enrichment Analysis (ssGSEA) demonstrated significantly increased enrichment of lysosomal lumen, lysosome organization, secondary lysosome, and autophagosome-lysosome fusion gene sets in the *Gpnmb*-high macrophage cluster relative to *Gpnmb*-low macrophage populations (**Fig. 3C**). Moreover, *Gpnmb* expression positively correlated with enrichment for genes annotated to lysosomal lumen within the *Gpnmb*-high cluster (**Fig. S6B**), further supporting a close association between GPNMB expression and lysosomal pathway activity.

Interestingly, the *Gpnmb*-associated transcriptional profile partially overlapped with disease-associated microglia (DAM) signatures described in neurodegeneration^22^. Injury-associated *Gpnmb^+^* macrophages exhibited elevated expression of genes linked to lysosomal activity, phagocytic remodeling, and DAM-like programs, including *Hexb, Ctss, Cd68, Ctsb, Trem2, Apoe, Lgals3, Cst7, Axl, and Tyrobp* (**Fig. 3D**). In addition, injury-associated *Gpnmb*⁺ macrophages expressed genes involved in myelin-derived lipid handling and cholesterol efflux, including *Lpl, Abca1, Abcg1,* and *Fabp5*, pathways that are essential for the processing and disposal of myelin-derived lipids^26,42,43^. Notably, *Lpl* and *Fabp5* were among the genes reduced in GPNMB-deficient macrophages, further supporting a role for GPNMB in lipid-processing adaptation. The coordinated enrichment of lysosomal, phagocytic, and lipid-processing genes suggests the emergence of a *Gpnmb*-positive macrophage state specialized for debris clearance and tissue remodeling during nerve regeneration. However, this transcriptional profile did not fully recapitulate the canonical DAM program, indicating that *Gpnmb*-expressing macrophages share key DAM-associated features while maintaining a distinct transcriptional profile.

Together, these analyses identify GPNMB as a marker of injury-associated macrophages enriched for lysosomal, lipid-processing, and reparative programs. The molecular signature of these cells is consistent with the defects observed in *Gpnmb*-deficient nerves and suggests that GPNMB is associated with macrophage states specialized for myelin processing, tissue remodeling, and successful nerve repair.

### GPNMB coordinates a lysosomal adaptation program in iPSC-derived macrophages

To determine whether GPNMB regulates macrophage transcriptional responses associated with lysosomal function, we performed bulk RNA sequencing of GPNMB knockout (KO) and non-targeting control (NT11) iPSC-derived macrophages exposed to myelin debris. Bafilomycin A1 (BafA1) and HBSS were included as reference perturbations to induce lysosomal stress and nutrient starvation-mediated autophagy, respectively. Principal component analysis demonstrated robust clustering of biological replicates and segregation by treatment condition, with additional separation between NT11 and GPNMB KO samples observed in PC3 and PC4 (**Fig. S7A**). Bafilomycin A1 (BafA1) and HBSS were included as reference perturbations to induce lysosomal stress and nutrient starvation–mediated autophagy, respectively. Gene ontology-based enrichment analysis revealed widespread transcriptional alterations in GPNMB-deficient macrophages across experimental conditions and time points (**Fig. 4A**). Processes and cellular components related to endosomal and lysosomal compartments, autophagy, vesicle trafficking, mitochondrial function, and immune responses were consistently altered in GPNMB KO cells, indicating broad disruption of pathways required for macrophage adaptation to myelin and lysosomal stress. Consistent with these pathway-level alterations, differential expression analysis identified numerous genes involved in lysosomal function, lipid handling, and immune responses among the most significantly altered transcripts in GPNMB-deficient macrophages across treatment conditions and time points (**Fig. S7B**).

**Figure 4.**
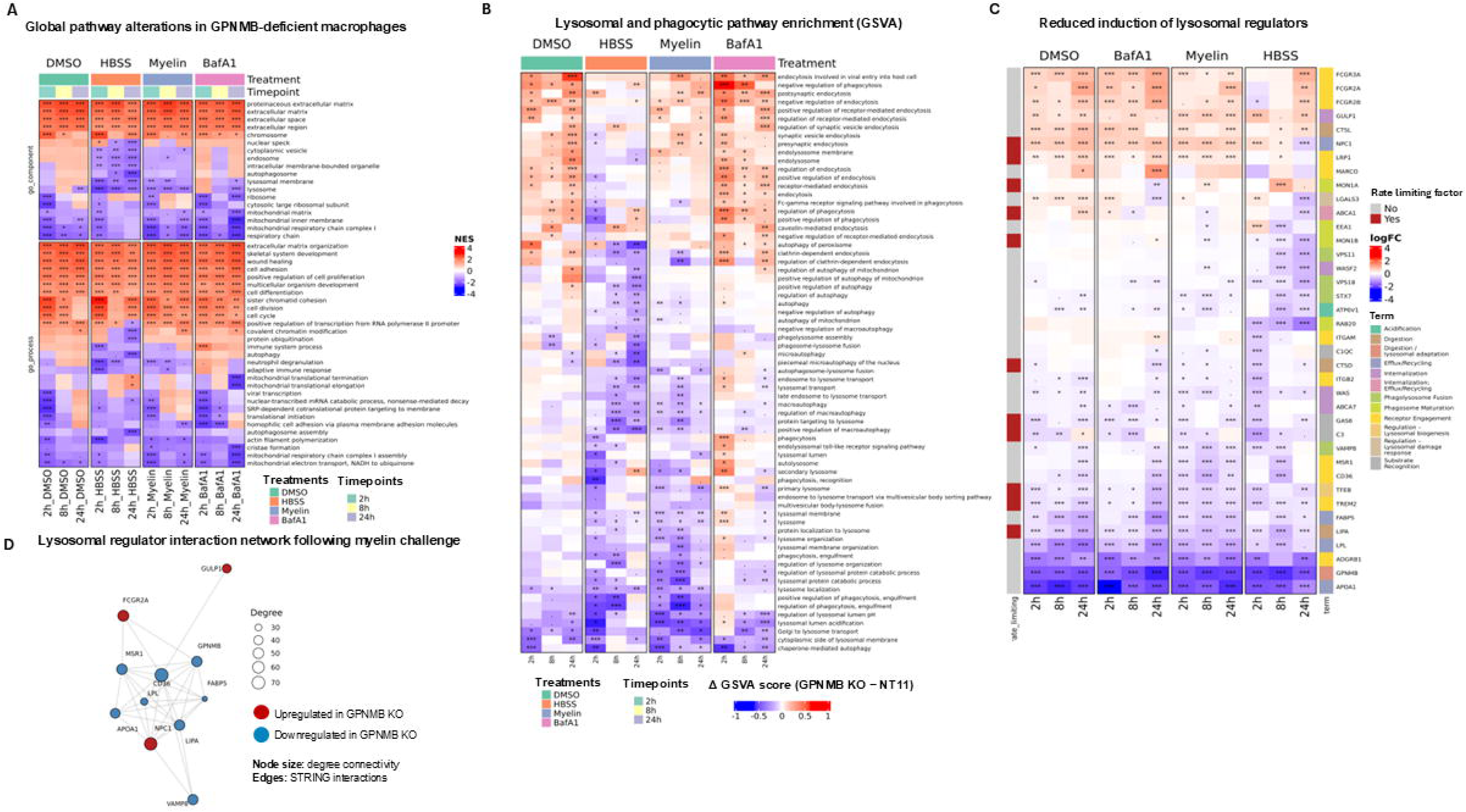
GPNMB deficiency suppresses lysosomal and phagocytic transcriptional programs in macrophages. **(A)** Heatmap showing normalized enrichment scores (NES) derived from unbiased gene set enrichment analysis (GSEA) of genes differentially expressed in GPNMB knockout (KO) relative to non-targeting control (NT11) macrophages following treatment with DMSO, HBSS, myelin, or bafilomycin A1 (BafA1) for 2, 8, or 24 h. Selected significantly enriched Gene Ontology (GO) biological process and cellular component terms are shown. Color scale indicates pathway enrichment (red, positive NES; blue, negative NES). Asterisks denote pathway significance based on Benjamini-Hochberg (BH)-adjusted p values: *** p ≤ 0.001, ** p ≤ 0.01, * p ≤ 0.05, . p ≤ 0.1, no symbol p > 0.1. **(B)** Heatmap showing differences in GSVA enrichment scores (ΔGSVA; GPNMB KO minus NT11) for selected phagocytosis-and lysosome-related gene sets across treatment conditions and time points. **(C)** Heatmap showing relative changes in mRNA abundance (logFC) of a manually curated set of lysosomal regulators in GPNMB knockout (KO) relative to non-targeting control (NT11) iPSC-derived macrophages following treatment with DMSO, bafilomycin A1 (BafA1), myelin, or HBSS for 2, 8, or 24 h. Only genes significantly regulated in at least one condition are shown (see Table SX for the full gene list). Genes were annotated according to their reported roles in lysosomal biology and whether they have been described as rate-limiting regulators of lysosomal function. Left annotations indicate rate-limiting regulators, whereas right annotations indicate the primary lysosomal process associated with each gene. Color scale represents logFC (GPNMB KO versus NT11), with red indicating higher and blue indicating lower expression in GPNMB KO macrophages. Asterisks indicate significantly differentially expressed genes (adjusted *P* < 0.05). **(D)** STRING protein-protein interaction network showing proteins encoded by genes differentially expressed in GPNMB KO macrophages following myelin treatment. Proteins encoded by genes downregulated in GPNMB KO macrophages form a highly interconnected network. Node color indicates differential expression relative to NT11 controls (red, upregulated; blue, downregulated). Node size reflects degree connectivity within the network.

To specifically assess pathways related to lysosomal function and cargo processing, we performed gene set variation analysis (GSVA) using lysosomal and phagocytic gene signatures. Across treatments, GPNMB-deficient macrophages exhibited reduced enrichment of pathways involved in phagocytosis, receptor-mediated endocytosis, lysosomal transport, autophagosome maturation, autophagosome-lysosome fusion, lysosome organization, and lysosomal acidification relative to control cells (**Fig. 4B**). These effects were particularly pronounced following myelin exposure and lysosomal stress induction, suggesting that GPNMB facilitates the transcriptional adaptation required for the uptake and processing of myelin-derived cargo and maintenance of lysosomal homeostasis.

To identify candidate regulators underlying these pathway-level changes, we examined the expression of genes associated with lysosomal function, lipid handling, membrane trafficking, and phagocytic activity. Multiple genes involved in cargo recognition and phagocytic uptake were reduced in GPNMB-deficient macrophages, including the scavenger receptors *CD36* and *MSR1*, which contribute to lipid and myelin debris uptake, and *MARCO*^44,45,46,47^, which participates in scavenger receptor-mediated recognition and uptake of extracellular cargo, as well as *FCGR2A* and *FCGR2B*, Fcγ receptors that regulate phagocytic and endocytic functions (**Fig. 4C**). Likewise, several genes implicated in lysosomal degradation and trafficking, including *LIPA*, *CTSD*, *CTSL*, *VAMP8*, and *ATP6V0A1*, exhibited reduced expression levels, consistent with impaired lysosomal adaptation. Additional decreases were observed in genes involved in lipid metabolism and macrophage activation, including *LPL*, *FABP5*, and *APOA1*. Notably, *LPL* and *FABP5* were enriched in GPNMB-high injury-associated macrophages in the iSNAT dataset (**Fig. 3A-D**) and were consistently reduced following GPNMB deletion in iPSC-derived macrophages. The concordance between the in vivo observational and in vitro loss-of-function datasets strengthens the evidence that GPNMB contributes to the maintenance of macrophage lipid-processing programs and supports a functional relationship between GPNMB expression and the regulation of *LPL* and *FABP5*.

Furthermore, expression of the lysosomal regulator *TFEB* and the microglial/macrophage activation marker *TREM2* was diminished. In contrast, *NPC1*, a key mediator of lysosomal cholesterol export, was increased in GPNMB-deficient cells, potentially reflecting a compensatory response to disrupted intracellular lipid handling. Together, these transcriptional changes indicate impaired activation of coordinated programs required for the recognition, uptake, and lysosomal processing of myelin-derived cargo, supporting a role for GPNMB in macrophage lysosomal adaptation.

Network analysis of differentially regulated lysosomal regulators further revealed an interconnected functional module centered on proteins involved in lipid uptake, lysosomal trafficking, and degradative capacity (**Fig. 4D**). Key regulators affected by GPNMB loss, including *CD36*, *MSR1*, *LIPA*, *VAMP8*, *LPL*, *FABP5*, and *NPC1*, occupied connected positions within the interaction network, highlighting coordinated regulation of pathways governing phagocytosis and lysosomal adaptation. To identify potential upstream regulators of these transcriptional changes, we inferred transcription factor activity from gene-expression changes in GPNMB KO iPSC-derived macrophages following myelin treatment. Several transcription factors exhibited genotype-dependent changes in predicted activity, many of which were not fully explained by changes in transcript abundance, suggesting altered regulatory network activity in GPNMB-deficient macrophages (**Fig. S7C**). MITF was among the top transcription factors predicted to exhibit reduced activity in GPNMB KO cells, and *MITF* expression was moderately decreased after 8 h of myelin exposure. Conversely, reanalysis of the injured sciatic nerve atlas (iSNAT) dataset revealed increased MITF activity in peripheral nerve macrophages during the recovery phase (**Fig. S7C**), coinciding with elevated *Gpnmb* expression (**Fig. 3B**). Given the established role of MITF in regulating lysosomal biogenesis, autophagy, and the expression of multiple lysosomal genes^48^, these complementary findings suggest that diminished MITF activity may contribute to the GPNMB-dependent repression of lysosomal and phagocytic programs.

Collectively, these findings demonstrate that GPNMB coordinates a lysosomal adaptation program that supports the recognition, uptake, and processing of myelin-derived cargo and promotes macrophage adaptation to myelin-associated and lysosomal stress. The concordant reduction of lysosomal and phagocytic gene programs in GPNMB-deficient macrophages, together with decreased predicted MITF activity, suggests that GPNMB may support these adaptive responses, at least in part, through a MITF-associated transcriptional network.

### GPNMB loss impairs autophagy-lysosome homeostasis in human iPSC-derived macrophages

To mechanistically substantiate our transcriptomic findings, we investigated whether GPNMB functionally regulates autophagy-lysosome activity in iPSC-derived macrophages. Under basal conditions, protein levels of the autophagy markers p62/SQSTM1 and LC3A/B were not significantly different between control and GPNMB KO macrophages (**Fig. S8B-D**). However, to assess autophagy-lysosome pathway responsiveness, macrophages were exposed to lysosomal inhibition or nutrient deprivation for 24 hours. Treatment with bafilomycin A1 (BafA1), which blocks lysosomal acidification and autophagosome-lysosome fusion, induced accumulation of p62/SQSTM1 in both genotypes, consistent with ongoing autophagic flux (**Fig. S8B, C**). LC3-II levels also increased following BafA1 treatment but accumulated to a lesser extent in GPNMB KO macrophages than in control cells (**Fig. S8B, D**). Likewise, treatment with AZD8055, a potent mTORC1/2 inhibitor that induces autophagy, significantly reduced LC3-II/I accumulation in GPNMB KO cells compared with NTC controls, whereas p62 levels showed only a modest, non-significant decrease (**Fig. S8B-D**). These data indicate that GPNMB contributes to the adaptive response of the autophagy-lysosome system under conditions of lysosomal or nutrient stress rather than regulating basal levels of autophagy-associated proteins.

Given that genotype-dependent differences emerged primarily under conditions that challenge the autophagy-lysosome system, we next investigated whether these effects were associated with altered upstream nutrient-sensing pathways. As mTORC1 coordinates lysosomal homeostasis and autophagic responses to metabolic stress, we examined mTORC1 pathway activity in GPNMB-deficient macrophages. Analysis of mTORC1 signaling revealed selective changes in pathway activity in GPNMB KO macrophages. Basal phosphorylation of S6RP was modestly increased in GPNMB KO cells compared with controls, whereas treatment with bafilomycin A1, AZD8055, or HBSS reduced p-S6RP levels in both genotypes (**Fig. S8E-F**). In contrast, phosphorylation of 4E-BP1 showed broadly comparable responses across conditions, and total 4E-BP1 protein levels remained largely unchanged (**Fig. S8E**). In addition, total TFEB abundance and phosphorylation status were broadly comparable between genotypes across treatment conditions (**Fig. S8H-I**). Although prolonged nutrient deprivation (24 h HBSS) reduced total protein abundance in both genotypes, consistent with severe cellular stress, no major genotype-dependent differences in TFEB regulation were observed at the protein level. These findings suggest that the impaired autophagic response observed in GPNMB-deficient macrophages is not explained by a gross disruption of TFEB protein abundance or phosphorylation, despite evidence for altered lysosomal-autophagic programs in the transcriptomic analyses.

We next asked whether GPNMB itself responds to perturbations of the autophagy-lysosome pathway. Bafilomycin A1 treatment markedly increased GPNMB protein abundance and induced a pronounced shift in electrophoretic mobility (**Fig. S8J-K**), consistent with altered post-translational processing. Chloroquine and AZD8055 also increased GPNMB levels, whereas HBSS treatment had only a modest effect. These findings indicate that GPNMB is responsive to lysosomal stress and support its association with autophagy-lysosome homeostasis, consistent with recent studies identifying GPNMB as both a marker and mediator of lysosomal stress adaptation in myeloid cells^49,17^.

Although the molecular basis of the mobility shift remains to be determined, these findings are consistent with the emerging view that GPNMB functions as part of an adaptive response to lysosomal stress^17,50,51^. The selective alterations in autophagy-related protein responses observed in GPNMB-deficient macrophages, together with the stress-induced upregulation of GPNMB, support a role for GPNMB in cellular adaptation to lysosomal and autophagic stress. These findings provide a potential framework for understanding how loss of GPNMB may contribute to defective myelin-debris processing after nerve injury.

### GPNMB loss alters p62 organization and lysosomal association in human iPSC-derived macrophages

To further investigate whether GPNMB deficiency alters handling of autophagy-associated cargo, we examined the selective autophagy receptor p62/SQSTM1 together with the lysosomal marker LAMP1 by immunofluorescence microscopy. Under basal conditions, p62/SQSTM1 exhibited a predominantly diffuse cytoplasmic distribution in control macrophages, whereas GPNMB KO macrophages displayed a more punctate staining pattern (**Fig. 5A-B**). Although quantitative analysis did not reveal a significant difference in p62-positive area at baseline, GPNMB-deficient macrophages consistently exhibited higher levels of p62-positive structures across treatment conditions. This trend was most pronounced following lysosomal inhibition with bafilomycin A1, where p62 accumulation was significantly greater in GPNMB KO macrophages than in control cells (**Fig. 5A-C**). In contrast, treatment with AZD8055 markedly reduced p62 puncta abundance in both genotypes. Nutrient deprivation (HBSS) also reduced p62-positive structures in both genotypes. Nevertheless, GPNMB KO macrophages retained a more punctate p62 staining pattern and exhibited a trend toward higher p62 puncta abundance relative to control cells (**Fig. 5A-C**). A similar pattern of altered p62 distribution was observed in human iPSC-derived microglia, indicating that disruption of p62 handling is not restricted to macrophages (**Fig. S9F**). Beyond alterations in p62-positive structures, GPNMB-deficient macrophages exhibited a pronounced lysosomal defect characterized by significantly reduced LAMP1 immunoreactivity (**Fig. 5A, D**) and fewer LAMP1-positive lysosomal puncta (**Fig. 5A, E**), indicating a reduction in lysosomal compartment abundance and/or integrity. Consistent with this phenotype, colocalization of p62-positive structures with LAMP1-positive lysosomes was significantly reduced in GPNMB KO macrophages (**Fig. 5B, F**), suggesting impaired coupling of autophagic cargo to lysosomal compartments. In control macrophages, GPNMB exhibited extensive colocalization with LAMP1-positive lysosomes (**Fig. 5A, G**), supporting a close spatial association between GPNMB and the lysosomal compartment (**Fig. 5A**; high-magnification images in **Fig. 5B, G**). Together, these findings indicate that GPNMB is required to maintain lysosomal homeostasis and organization in macrophages, with its loss leading to reduced lysosomal abundance and impaired association of p62-positive cargo with lysosomal compartments.

**Figure 5.**
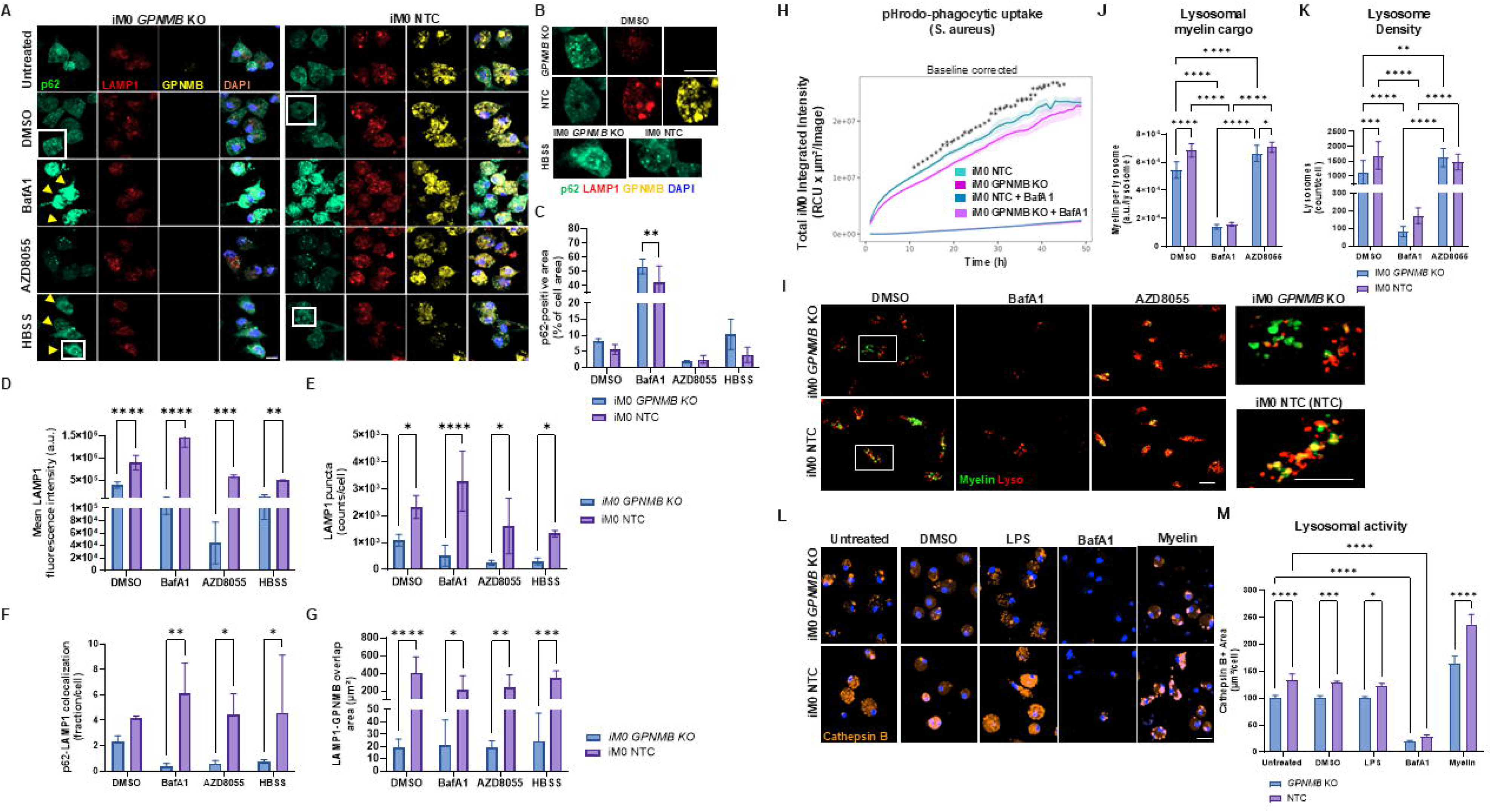
GPNMB deficiency impairs lysosomal function and autophagic cargo processing in human iPSC-derived macrophages. **(A)** Representative confocal images of NTC and GPNMB KO human iPSC-derived macrophages treated with DMSO, bafilomycin A1 (BafA1), AZD8055, or HBSS and stained for p62/SQSTM1 (green), LAMP1 (red), GPNMB (yellow), and DAPI (blue). Arrowheads indicate p62-positive structures following BafA1 treatment. Insets show higher-magnification views of the boxed regions. Scale bars, 20 μm (main panels) and 10 μm (insets). **(B)** High-magnification representative confocal images highlighting p62/SQSTM1 (green), LAMP1 (red), and GPNMB (yellow) positive compartments in NTC and GPNMB KO macrophages following DMSO or HBSS treatment. Under starvation conditions (HBSS), NTC macrophages exhibited a predominantly diffuse p62 staining pattern, whereas GPNMB KO macrophages retained prominent p62-positive puncta and altered intracellular p62 distribution. Scale bar, 10 μm. **(C)** Quantification of p62-positive area, expressed as the percentage of total cell area, in GPNMB KO and NTC human iPSC-derived macrophages following 24 h treatment with DMSO, BafA1, AZD8055, or HBSS. **(D-E)** Quantification of mean LAMP1 fluorescence intensity per cell (**D**) and LAMP1-positive puncta per cell (**E**) in GPNMB KO and NTC human iPSC-derived macrophages following 24 h treatment with DMSO, BafA1, AZD8055, or HBSS. **(F–G)** Quantification of p62-LAMP1 colocalization, expressed as the fraction of p62 signal colocalizing with LAMP1-positive lysosomal compartments (**F**), and GPNMB-LAMP1 colocalization, expressed as the total overlap area between GPNMB- and LAMP1-positive structures (µm²) (**G**), in GPNMB KO and NTC human iPSC-derived macrophages following 24 h treatment with DMSO, BafA1, AZD8055, or HBSS. **(H)** Kinetics of phagocytosis in GPNMB KO and NTC iM0 macrophages assessed by Incucyte live-cell imaging following exposure to pHrodo™-labeled *Staphylococcus aureus* bioparticles. Phagocytic uptake was quantified as integrated red fluorescence intensity (RCU/μm²/image) at the indicated time points. Bafilomycin A1 (BafA1), an inhibitor of lysosomal acidification, was included as a positive control for impaired lysosomal processing of internalized cargo. Individual traces represent independent wells. **(I)** Representative confocal images showing uptake and lysosomal processing of pHrodo-labeled myelin (green) in NTC and GPNMB KO human iPSC-derived M0 macrophages (iM0) following treatment with DMSO, bafilomycin A1 (BafA1), or AZD8055. Lysosomes were labeled with LysoTracker Red (red). Bottom panels show higher-magnification views of the boxed regions. Scale bars, 40 μm (main panels) and 10 μm (high-magnification insets). **(J)** Quantification of myelin-associated lysosomal signal (A.U.), defined as the association of pHrodo-labeled myelin (green) with LysoTracker-positive lysosomal compartments (red), in NTC and GPNMB KO macrophages following the indicated treatments. Data are presented as mean ± SD. Statistical analysis was performed using two-way ANOVA followed by Tukey’s multiple-comparisons test. **(K)** Quantification of LysoTracker-positive lysosomes per cell in NTC and GPNMB KO human iPSC-derived M0 following treatment with DMSO, BafA1, or AZD8055 in the presence of pHrodo-labeled myelin. Data are presented as mean ± SD. Statistical analysis was performed using two-way ANOVA followed by Tukey’s multiple-comparisons test. **(L)** Representative confocal images showing Cathepsin B activity detected using the Magic Red Cathepsin B reporter (orange) in NTC and GPNMB KO human iPSC-derived M0 macrophages (iM0) following treatment with DMSO, LPS, bafilomycin A1 (BafA1), or myelin. Nuclei were counterstained with DAPI (blue). Scale bar, 20 μm. **(M)** Quantification of lysosomal proteolytic activity in GPNMB KO and NTC macrophages under untreated conditions or following treatment with DMSO, LPS, bafilomycin A1 (BafA1), CA-074, or myelin. Cathepsin B activity was measured using the Magic Red Cathepsin B reporter assay and quantified as Magic Red-positive area per cell 2 h after probe addition. Data are presented as mean ± SD. Statistical significance was assessed using two-way ANOVA followed by Tukey’s multiple comparisons test. Data are presented as mean ± SD. Statistical significance was assessed using two-way ANOVA followed by Tukey’s multiple-comparisons test or Šidák’s multiple-comparisons test. Significance is indicated on the graphs. *p* < 0.05, \**p* < 0.01, \*\**p* < 0.001, \*\*\**p* < 0.0001, \*\*\*\**p* < 0.0001.

### GPNMB is required for lysosomal adaptation to myelin-induced stress

Because persistent myelin debris accumulation in vivo suggested defective cargo processing, we first assessed phagocytic activity in human iPSC-derived macrophages using pHrodo-labeled *Staphylococcus aureus* bioparticles. Live-cell imaging over a 24 h time course revealed lower pHrodo signal accumulation in *GPNMB* KO macrophages compared with control cells, indicating reduced phagocytic activity and/or impaired phagosome acidification (**Fig. 5H; Fig. S9A**).

To determine whether these defects extended to a physiologically relevant substrate, we next challenged macrophages with pHrodo-labeled myelin for 24 h and performed endpoint imaging with LysoTracker to assess the association of internalized myelin with acidic lysosomal compartments and to quantify lysosome abundance. Under vehicle-treated conditions (DMSO), *GPNMB* KO macrophages exhibited reduced myelin-associated lysosomal signal and decreased lysosome abundance compared with WT (NTC) cells (**Fig. 5I-K; Fig. S9B-C**). As expected, bafilomycin A1 markedly reduced both myelin-associated lysosomal signal and lysosome abundance in both genotypes (**Fig. 5I-K; Fig. S9B-C**). In contrast, treatment with the mTOR inhibitor AZD8055 restored myelin-associated lysosomal signal and lysosome abundance in *GPNMB* KO macrophages (**Fig. 5I-K; Fig. S9B-C**). These findings show that GPNMB contributes to efficient lysosomal processing of myelin-derived cargo. The partial restoration of lysosome abundance and myelin-associated lysosomal signal by AZD8055 further indicates that defects associated with GPNMB deficiency may remain amenable to pharmacological modulation of lysosome-autophagy pathways.

To directly assess lysosomal degradative capacity, we next quantified Cathepsin B activity using the Magic Red reporter assay (**Fig. 5L**). Under basal untreated and DMSO-treated conditions, *GPNMB* KO macrophages exhibited reduced Cathepsin B-positive area compared with control cells, consistent with diminished steady-state lysosomal proteolytic activity (**Fig. 5L-M, S9D)**. LPS stimulation did not reveal major genotype-dependent differences, whereas bafilomycin A1 reduced Cathepsin B activity in both groups, as expected from its inhibition of lysosomal acidification. In contrast, myelin challenge triggered a robust increase in Cathepsin B activity in control macrophages but elicited a substantially weaker response in *GPNMB* KO cells (**Fig. 5L-M, S9D**), suggesting an impaired ability to upregulate lysosomal degradative function in response to increased myelin burden. A similar reduction in lysosomal proteolytic activity was observed in GPNMB-deficient human iPSC-derived microglia (**Fig. S9E**), supporting a broader role for GPNMB in maintaining lysosomal function across human myeloid cell types.

Collectively, these results demonstrate that loss of GPNMB disrupts multiple aspects of lysosomal biology, including lysosome abundance, degradative activity, and maintenance of LAMP1-positive lysosomal organization. These defects impair processing of myelin-derived cargo and limit the ability of macrophages to mount an effective lysosomal response to increased degradative demand. Because efficient lysosomal function is required for the clearance of autophagic cargo, we next investigated whether GPNMB deficiency also affects p62/SQSTM1 turnover and autophagy-associated cargo processing.

### GPNMB-deficient macrophages are selectively vulnerable to phagocytic and lysosomal stress

Finally, we asked whether the lysosomal defects associated with GPNMB deficiency compromise macrophage viability under stress. Because sustained lysosomal dysfunction can promote apoptotic cell death through defective cargo clearance and cellular stress^52–54^, we examined apoptotic responses in GPNMB-deficient macrophages under conditions that challenge lysosomal homeostasis. iPSC-derived macrophages were exposed for 24 h to DMSO, bafilomycin A1, AZD8055, myelin, or camptothecin (positive control for apoptosis; induces DNA damage through topoisomerase I inhibition), and apoptosis was assessed using Annexin V, caspase-3/7 activation, and Cytotox readouts.

Under basal conditions, control and *GPNMB* KO macrophages showed similarly low levels of apoptosis, indicating that loss of GPNMB does not induce spontaneous cell death. In contrast, myelin exposure induced a pronounced apoptotic response in *GPNMB* KO macrophages, characterized by increased Annexin V signal, caspase-3/7 activation, and Cytotox positivity. Control macrophages showed limited apoptotic activation under the same conditions (**Fig. 6A-C**). Bafilomycin A1 further increased Annexin V-positive cells in *Gpnmb* KO macrophages, indicating enhanced early apoptotic responses to lysosomal stress. In contrast, camptothecin induced apoptosis in both genotypes with only modest differences. These data indicate that *Gpnmb*-deficient macrophages are selectively vulnerable to lysosomal and phagocytic stress rather than broadly hypersensitive to all apoptotic stimuli.

**Figure 6:**
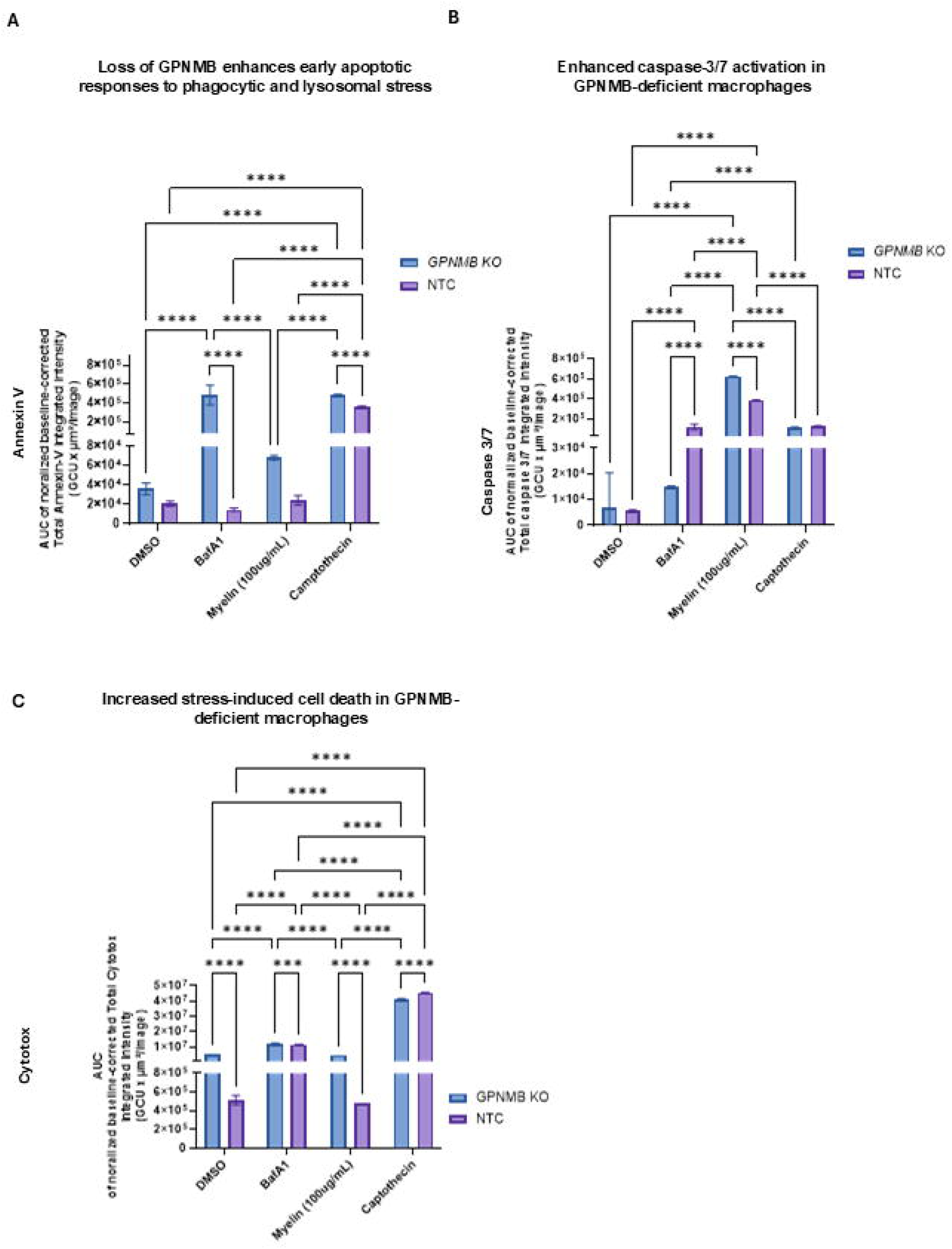
GPNMB protects macrophages from lysosomal and phagocytic stress-induced apoptosis. **(A-C)** Apoptotic responses of NTC and *GPNMB* KO human iPSC-derived macrophages following 24 h treatment with DMSO, myelin, bafilomycin A1 (BafA1), or camptothecin. Apoptosis was assessed by Annexin V (**A**), caspase-3/7 activation (**B**), and Cytotox (**C**) readouts. Signals were normalized to cell number at each time point and quantified as area under the curve (AUC). Data are presented as mean ± SD. Statistical analysis was performed using two-way ANOVA followed by Tukey’s multiple-comparisons test.

Together, these findings indicate that GPNMB protects macrophages from the deleterious consequences of sustained phagocytic and lysosomal stress. Consistent with the defects in lysosomal fitness, cargo processing, and autophagy-lysosome alterations described above, GPNMB-deficient macrophages exhibit increased susceptibility to apoptosis when challenged with myelin burden or lysosomal dysfunction.

## Discussion

Peripheral nerve regeneration requires coordinated immune and glial responses to remove inhibitory debris and restore axonal connectivity, yet the molecular mechanisms linking these processes are incompletely understood. Efficient regeneration depends on macrophage-mediated debris clearance and Schwann cell reprogramming during Wallerian degeneration, which together govern axon regrowth and remyelination^30,55^. Here, we identify GPNMB as a key regulator operating across macrophages and spinal cord microglia that couples myelin debris clearance with the acquisition of pro-repair cellular states. By integrating in vivo and in vitro approaches, we show that GPNMB is required for lysosome-dependent reparative programs in injury-responsive myeloid cells that drive efficient tissue remodeling and nerve regeneration.

In rodents, *Gpnmb* is rapidly induced following peripheral nerve injury and remains elevated throughout both the degenerative and regenerative phases. This sustained expression contrasts with the transient inflammatory response that follows injury and is instead consistent with a repair-associated program. Previous work demonstrated that administration of soluble GPNMB promotes peripheral nerve regeneration following sciatic nerve crush injury, supporting a beneficial role for GPNMB in nerve repair^30^. The beneficial effects of soluble GPNMB administration reported previously and the regenerative defects observed here in GPNMB-deficient animals are consistent with a model in which both endogenous and exogenous GPNMB enhance repair-associated cellular programs. Our findings extend this concept by demonstrating that endogenous GPNMB is required for normal functional recovery following nerve injury. Loss of GPNMB resulted in persistent abnormalities detected by MRI, defective myelin clearance, impaired lysosomal adaptation, and reduced acquisition of repair-associated cellular states across multiple injury-responsive cell populations. By combining in vivo and in vitro analyses, we identify myelin processing and lysosome-dependent repair programs as key biological processes regulated by GPNMB during regeneration. Together, these findings support an important cell-intrinsic role for GPNMB in coordinating lysosome-dependent repair programs required for efficient nerve regeneration, whereas previous studies primarily focused on the effects of exogenous soluble GPNMB.

GPNMB is expressed by both macrophages and Schwann cells, supporting the possibility of a coordinated injury-induced repair program rather than a lineage-restricted function. In the injured peripheral nerve, its expression coincides with the reprogramming of Schwann cells and resident macrophages toward regenerative states^26,30^. Consistent with previous reports, our data show that GPNMB is enriched in macrophage populations associated with lipid metabolism, lysosomal function, and tissue repair^9,29^. Reanalysis of single-cell transcriptomic datasets further revealed that *Gpnmb*-high macrophages are enriched for gene programs associated with lysosome organization, lysosomal lumen function, secondary lysosome formation, and autophagosome-lysosome fusion. Moreover, *Gpnmb* expression positively correlated with lysosomal pathway activity across injury-associated macrophage populations, supporting a close relationship between GPNMB expression and the acquisition of a reparative macrophage state specialized for lysosomal adaptation.

This lysosomal signature was mirrored in human iPSC-derived macrophages, where loss of GPNMB resulted in broad suppression of lysosomal and phagocytic transcriptional programs. Bulk RNA sequencing demonstrated reduced enrichment of pathways involved in phagocytosis, lysosomal organization, and autophagosome-lysosome function, accompanied by altered expression of key regulators of cargo uptake, lysosomal trafficking, and lipid processing, including CD36, MSR1, LIPA, VAMP8, FABP5, and APOA1. Together with our single-cell analyses showing that *Gpnmb*-high macrophages are enriched for lysosomal and autophagosome-lysosome fusion programs, these findings suggest that GPNMB contributes not only to lysosomal activity at a functional level but also to the transcriptional networks that enable macrophage adaptation to degradative and lipid-associated stress. Consistent with this notion, transcription factor activity analysis identified MITF as one of the transcription factors most strongly affected by GPNMB deficiency in myelin-challenged macrophages^56^. Moreover, MITF activity was similarly increased in macrophages from injured peripheral nerve, suggesting that transcriptional programs associated with lysosomal adaptation are shared between the in vitro myelin challenge model and the in vivo regenerative response. Given the established role of MITF family transcription factors in regulating lysosomal biogenesis, autophagy, and cellular adaptation to metabolic stress^57,58^, these findings further link GPNMB to transcriptional networks governing lysosomal adaptation and degradative function in injury-responsive macrophages. Importantly, disruption of this lysosome-centered program in *Gpnmb*-deficient mice was associated with impaired acquisition of repair-associated macrophage signatures, defective debris clearance, and delayed functional recovery following nerve injury.

Efficient myelin clearance is a critical determinant of successful peripheral nerve regeneration, as persistence of myelin-derived debris can inhibit axonal regrowth and delay repair. Macrophages coordinate this process through phagocytic uptake and lysosomal degradation of damaged myelin, and disruption of lysosomal or autophagy-associated pathways has been linked to impaired regenerative outcomes following nerve injury^59,60^. Our findings position GPNMB within this broader network of regulators by identifying a role in supporting the lysosomal and phagocytic programs required for adaptation to myelin burden. The observed defects in lysosomal function, myelin processing, and acquisition of repair-associated macrophage states in GPNMB-deficient cells suggest that GPNMB contributes to the degradative machinery necessary for efficient tissue remodeling during nerve regeneration.

In vivo, GPNMB-deficient macrophages exhibited reduced CD68 expression and smaller size, suggesting impaired acquisition of a degradative injury-associated phenotype. Likewise, microglia showed reduced soma enlargement and failed to efficiently accumulate around injured motor neurons, indicating an attenuated injury response^61–64^. These alterations were associated with persistent peripheral nerve myelin debris, delayed functional recovery, and altered MRI profiles. Elevated magnetization transfer ratio (MTR) in GPNMB-deficient nerves is more consistent with the accumulation of macromolecular debris than with productive remyelination, supporting a predominant defect in degradative processing and clearance rather than in debris uptake^65,66^.

Mechanistically, human iPSC-derived macrophages and microglia lacking GPNMB recapitulated key aspects of the in vivo phenotype. Although GPNMB deficiency modestly reduced myelin uptake and phagocytic activity, defects became more apparent at the level of lysosomal adaptation and cargo processing. GPNMB-deficient cells displayed reduced LAMP1 immunoreactivity and lysosomal puncta abundance, diminished cathepsin activity, impaired myelin-lysosome association, accumulation of p62-positive structures following lysosomal stress, and reduced association of p62-positive structures with lysosomal compartments.

Notably, autophagy initiation appeared largely preserved, whereas defects became increasingly evident under conditions of sustained cargo burden, suggesting an impaired capacity to maintain lysosomal function during degradative stress. Interestingly, lysosomal inhibition by bafilomycin A1 markedly increased GPNMB abundance and altered its electrophoretic mobility, suggesting stress-dependent post-translational regulation^17,18,67^. Consequently, myelin challenge precipitated a progressive loss of cellular fitness and increased susceptibility to apoptosis. GPNMB-deficient macrophages were not broadly hypersensitive to apoptosis, as camptothecin induced comparable responses in both genotypes. Instead, they displayed selective vulnerability to myelin burden and lysosomal stress, consistent with a primary defect in stress adaptation rather than basal cell survival. Together, these findings position GPNMB as a regulator of lysosomal adaptation that enables macrophages and microglia to cope with the metabolic and degradative demands imposed by myelin processing^18,19,49,67–69^.

Our findings are consistent with previous genetic evidence identifying *Gpnmb* as a modifier of macrophage lysosome function. Robinet et al., 2021^70^ showed that loss of functional *Gpnmb* reduces lysosomal degradative capacity in mouse macrophages and that restoration of wild-type *Gpnmb* rescues this defect. Notably, the p62 phenotype differed between cultured macrophages and injured nerve tissue: in vitro, GPNMB KO macrophages showed redistribution of p62 into punctate structures together with altered p62-LAMP1 association, whereas in vivo, *Gpnmb*-deficient nerves showed increased endoneurial p62 but reduced macrophage-associated p62 signal. This apparent discrepancy likely reflects the distinction between tissue-level p62 accumulation measured in vivo and the intracellular distribution of p62 within individual macrophages assessed in vitro, as well as differences in the cellular environment between these experimental systems.

Importantly, our findings extend beyond the peripheral nerve. Following sciatic nerve injury, spinal cord microglia upregulated GPNMB and acquired features associated with lysosomal activation and tissue remodeling. This aligns with studies in neurodegeneration identifying GPNMB as a key marker of disease-associated microglia (DAM), characterized by enhanced lysosomal activity and attenuated inflammatory signaling^22,25^. In contrast, GPNMB deficiency attenuated microglial activation, reduced CD68 expression, limited soma enlargement, and impaired accumulation around injured motor neurons. Together with our transcriptomic analyses showing that Gpnmb-positive injury-associated macrophages acquire DAM-like lysosomal and lipid-processing programs without fully adopting a canonical DAM identity, these findings support the existence of conserved GPNMB-dependent repair programs across the PNS–CNS axis^71^. We therefore propose that GPNMB functions as a critical regulator of lysosomal adaptation in injury-responsive myeloid cells, coupling efficient cargo processing to the acquisition of reparative cellular states during regeneration.

## Limitations and future directions

Several limitations should be considered. First, the use of constitutive *Gpnmb* null mice does not exclude potential developmental effects that may influence injury responses. Because GPNMB is expressed by both macrophages and Schwann cells following nerve injury, future lineage-specific loss-of-function studies will be required to determine the relative contribution of myeloid versus Schwann cell GPNMB to lysosomal adaptation, myelin clearance, and regenerative repair.

Second, although MTR imaging provides a sensitive non-invasive measure of tissue composition, it represents an indirect readout of myelin dynamics. Histological analyses demonstrated accumulation of degraded MBP-positive material in *Gpnmb*-deficient nerves, supporting the interpretation that persistent MRI abnormalities reflect retained myelin debris rather than newly formed myelin. Future studies using ultrastructural approaches such as electron microscopy may further clarify the relationship between debris clearance and remyelination. Finally, although our findings identify impaired lysosomal adaptation as a central consequence of GPNMB deficiency, direct demonstration that restoration of GPNMB function rescues lysosomal competence and degradative capacity remains an important next step. Re-expression studies in GPNMB-deficient human iPSC-derived macrophages, together with pharmacological enhancement of lysosomal function, will be important for establishing a causal relationship between GPNMB, lysosomal competence, and regenerative function. Given the functional rescue of lysosomal phenotypes observed following pharmacological mTOR inhibition, future studies should determine whether therapeutic enhancement of lysosomal adaptation can restore myelin processing and improve regenerative outcomes in the absence of GPNMB.

## Therapeutic implications

Our findings identify lysosomal fitness as a key determinant of regenerative success and position GPNMB as a central regulator of this process. These results suggest that therapeutic strategies aimed at enhancing GPNMB function or improving lysosomal resilience may help promote tissue repair in conditions characterized by impaired debris clearance and chronic phagocyte dysfunction, including peripheral neuropathies and neurodegenerative disorders such as ALS. The existence of a soluble form of GPNMB, together with the feasibility of viral-mediated gene delivery, raises the possibility of both protein- and gene-based therapeutic approaches. Future studies will be required to determine whether restoration of GPNMB function is sufficient to rescue lysosomal competence and regenerative capacity in disease settings.

## Conclusion

Together, these findings establish GPNMB as a key regulator of lysosomal adaptation in injury-responsive myeloid cells and identify this pathway as a critical determinant of efficient myelin clearance and successful nerve regeneration.

## Supporting information

supp. table 3 in support of the iSNAT data in Fig. 3 and S6

supp. table 1 link to the Bulk RNA seq data in Fig. 4 and S7

supp. table 1 link to the iSNAT data in fig. 3 and S6

## Acknowledgments

We thank the CM In Vivo Sciences team for their outstanding support and contributions to the in vivo studies.

We are grateful to all members of the DARe Pharmacology group for their scientific discussions, advice, guidance, and continuous support throughout this project. We especially thank Marie Ronco, Nathalie Accart Gris, and Arno Doelemeyer for their valuable insights, encouragement, and assistance in facilitating access to research platforms and core facilities.

We thank Inga Galuba (Discovery Sciences) for assistance with image acquisition.

We are grateful to the DARe In Vitro group, particularly Eun Yeong Bergsdorf, Benjamin Jourde, and Fanning Zeng, as well as Isabelle Fruh and Mitchel Germain (Discovery Sciences), for technical assistance, reagent support, and valuable scientific discussions. We also thank Yannic Knoepfli and Yara Haefeli (Neuroscience) for their support with the Incucyte platform.

We acknowledge Discovery Sciences for providing the facilities, infrastructure, and research resources that enabled this work. We particularly thank Jasmin Haegele and Marc Altorfer (Analytical Genomics, Discovery Sciences) for their support with bulk RNA sequencing, including sample processing and library preparation.

Finally, we thank Ina Kramer for insightful scientific discussions, guidance, encouragement, and unwavering support throughout this project.

N.O.A. was supported by the Novartis Postdoctoral Fellowship Program.

## Author Contributions

**Conceptualization,** N.O.A., M.B.,; **Methodology,** N.O.A., M.P., M.O., G.S., G.K., M.M., Z.T., D.G., and I.C.; **Investigation,** N.O.A., M.P., M.O., T.D., I.O., C.P., L.K., D.C.-I., E.G., C.L., V.V.S., L.H., and C.M.; **Formal Analysis,** N.O.A., I.O., O.D., Z.T., D.G., I.C., and N.B.; **Data Curation,** I.O., O.D., and N.B.; **Resources,** M.M. and M.N.; **Writing – Original Draft,** N.O.A.; **Writing – Review & Editing,** M.L., D.S., N.B, O.D., M.N., M.B.,; **Supervision,** M.B.

## Methods

### In Vivo Experiments Animals

All animal procedures were performed in accordance with Swiss animal welfare legislation and were approved by the Kantonales Veterinäramt Basel-Stadt under animal experimentation license BS-2804 (“Identification of therapeutics to improve neuromuscular function following peripheral nerve injury in rodents”).

C57BL/6NRj mice (Charles River Laboratories, Germany) and DBA/2J mice carrying a spontaneous loss-of-function mutation in *Gpnmb* (Gpnmb^R150X; Jackson Laboratory strain #000671) together with isogenic DBA/2J-Gpnmb^+/SjJ controls (strain #007048) were used in this study.

C57BL/6 mice were used for temporal analyses of nerve regeneration and injury-induced *Gpnmb* expression (11-week-old at surgery; n = 24 total). DBA/2J mice were used to assess the functional consequences of GPNMB deficiency (8–10-week-old at surgery; n = 30 per genotype).

Animals were allowed to acclimatize for at least 7 days prior to experimentation. They were housed under standard conditions (temperature 20–24 °C, relative humidity ≥40%, 12 h light/dark cycle) in ventilated cages with a maximum of 4 animals per cage and provided with food and water ad libitum.

### Mouse sciatic nerve crush (SNC)

Sciatic nerve crush injury was performed as described previously (Giorgetti E. et al., 2019). Animals were anesthetized by induction with 4% isoflurane and maintained under continuous inhalation of 1.5–2% isoflurane on a heated surface.

The skin was incised posterior and parallel to the femur. The left sciatic nerve was exposed at mid-thigh level and crushed for 15 s proximal to its bifurcation using Dumont #5 forceps.

Powdered charcoal was applied to the forceps to mark the crush site. Following injury, the nerve was repositioned and the muscle was closed with absorbable sutures, and the skin incision was sealed with wound clips.

For sham-operated controls, the nerve was exposed but not crushed.

Post-operative analgesia was provided using non-steroidal anti-inflammatory drugs (Meloxicam, 2–5 mg/kg, subcutaneously), administered before surgery and continued for up to 48 h post-operatively.

Animals were allowed to recover from anesthesia and returned to their home cages.

### Magnetic resonance imaging (MRI)

MRI measurements were performed using a 7 T Biospec 70/16 spectrometer (Bruker Medical Systems, Ettlingen, Germany) operating with Paravision software.

Spontaneously breathing, isoflurane-anesthetized animals were positioned in prone position in a dedicated cradle, and body temperature was maintained at approximately 37 °C. Images of both hind limbs were acquired simultaneously but mice had to be repositioned for imaging the spinal cord. Respiration was monitored continuously during data acquisition. Details about acquisition parameters are provided as supplemental material.

### Sciatic nerve acquisition

A T2-weighted multislice RARE sequence was used for anatomical localization of the sciatic nerve. Multislice FLASH acquisitions were subsequently performed to determine magnetization transfer ratio (MTR) as a measure of tissue macromolecular content, including myelin. Details about acquisition parameters are furnished as supplemental materials.

MTR was calculated as:

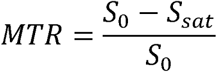

where *S*₀ and *Ssat* represent signal intensities obtained without and with magnetization transfer saturation. Regions of interest (ROIs) were defined on anatomical images and transferred to corresponding FLASH datasets for quantitative analysis.

### Spinal cord acquisition

For assessment of central responses to peripheral nerve injury, MRI datasets were acquired at the level of the spinal cord, focusing on lumbar (L1–L6) and thoracic segments.

Initial anatomical localization was performed using pilot scans followed by high-resolution coronal acquisitions covering the full rostrocaudal extent of the lumbar and thoracic spinal cord.

FLASH-based sequences were used to quantify MTR in defined spinal cord regions, using the same acquisition parameters as for peripheral nerve imaging except for the geometry. Measurements were performed with and without magnetization transfer saturation pulses to allow calculation of MTR values.

Regions of interest (ROIs) were manually defined on anatomical reference images for lumbar and thoracic segments and subsequently applied to MTR datasets for analysis.

### Data processing

MTR and T2 analyses were performed using Paravision software. ROIs defined on anatomical images were consistently applied across corresponding datasets. Signal intensities were quantified to compute MTR values and to assess tissue-specific changes over time.

### Electrophysiological assessments (CMAP)

Electrophysiological measurements were performed in isoflurane-anesthetized animals using an ADInstruments recording system.

The sciatic nerve was stimulated using monopolar needle electrodes placed in the gluteal region at the sciatic notch. Recordings were obtained from the gastrocnemius muscle using an active electrode, with a reference electrode placed in the hindlimb.

For CMAP measurements, the nerve was stimulated with single pulses (0.1–0.5 ms), and the current was gradually increased to achieve maximal amplitude responses.

Measurements were performed at baseline and at weekly intervals following nerve injury.

### Toe spread assessment

Motor function recovery was evaluated using the toe spread reflex. Mice were gently lifted by the tail and hindlimb toe spreading was visually assessed.

Responses were scored as follows:

- 0: no spreading
- 1: intermediate spreading
- 2: full spreading

Mean scores were calculated for each group at the respective time points.

### Compound muscle action potential (CMAP) analysis

CMAP amplitudes were analyzed using a linear mixed-effects model (LME) implemented in the lme4 package (R v4.6.0). The model included strain (*Gpnmb*-null or *Gpnmb*-WT), timepoint, and surgery type (SNC or sham) as fixed effects, together with all interaction terms, and a random intercept for each animal to account for repeated measurements:

CMAP ∼ Strain * Timepoint * Surgery + (1|Animal_ID)

Estimated marginal means (EMMs) were calculated using the emmeans package. Pairwise comparisons between strains within the SNC group were performed at each timepoint and resulting *p values* were corrected for multiple testing using the Holm method.

### Toe spread analysis

Toe-spread scores were assessed on a 0–2 ordinal scale and compared between the injured (ipsilateral) and uninjured (contralateral) hindlimbs of sciatic nerve crush (SNC) animals. For each strain (*Gpnmb*-WT and *Gpnmb*-null), toe-spread scores were compared at each post-injury timepoint using the Wilcoxon rank-sum test (Mann-Whitney U test). The median difference between ipsilateral and contralateral scores was calculated as a measure of functional deficit. P values were adjusted for multiple comparisons across all strain × timepoint combinations using the Holm method.

### Tissue collection and histology

At defined endpoints, animals were euthanized by overdose of isoflurane and transcardially perfused with ice-cold phosphate-buffered saline (PBS) supplemented with 5 mM EDTA, followed by 4% paraformaldehyde (PFA) at a constant flow rate of 5 mL/min.

Sciatic nerves (proximal and distal to the crush site) and lumbar spinal cord (L1–L6) were collected.

Tissues were post-fixed in 4% PFA overnight, cryoprotected in 30% sucrose, embedded in OCT, and stored for cryosectioning.

### Cryosectioning and immunofluorescence

Following perfusion and tissue processing, sciatic nerve and spinal cord samples were cryoprotected in 30% sucrose, embedded in OCT, and stored at −80 °C until sectioning.

Cryosectioning was performed using a Cryostat (Thermo Scientific CryoStar NX70) set between −18 °C and −23 °C. Tissue blocks were equilibrated overnight at -20°C and inside the cryostat for at least 1 h prior to sectioning to ensure optimal cutting quality and preservation of tissue integrity.

### Sciatic nerve sections

Sciatic nerves were dissected relative to the charcoal-marked crush site, allowing separation of proximal and distal regions and preparation of both longitudinal and transverse sections.

Sciatic nerves were sectioned at 12 µm thickness and mounted onto HistoBond-coated glass slides (HistoBond®+M; Marienfeld Superior). Sections were air-dried overnight at room temperature to ensure stable adhesion prior to staining.

Longitudinal or transverse orientations were selected depending on the experimental time point and region relative to the crush site.

When required antigen retrieval was performed using sodium citrate buffer (10 mM, pH 8.0) at 80 °C (water bath) for 45 min.

Immunostaining was performed using the same procedure as for free-floating sections, including blocking, primary antibody incubation (24h), washing, and secondary antibody labeling.

### Spinal cord sections (free-floating)

Lumbar spinal cord sections (L1–L6) were cut at 40 µm thickness, collected serially, and washed in 0.1 M PBS (w/o Ca^2+^/Mg^2+^).

When required, antigen retrieval was performed by incubating the sections in 10 mM sodium citrate buffer (pH 8.0) at 80 °C for 45 min in a water bath.

Sections were then blocked in 10% donkey serum (or 3% BSA) containing 0.3% Triton X-100 for 2 h at room temperature.

Primary antibodies diluted in blocking buffer were applied for 48 h at 4 °C on a shaker. Following incubation, sections were washed three times (30 min each) in 0.1 M PBS and incubated with appropriate fluorophore-conjugated secondary antibodies for 2 h at room temperature.

After secondary staining, sections were washed (3 × 30 min) and mounted using ProLong Gold antifade mounting medium (Thermo Fisher Scientific).

Tissue collections, cryosectioning and immunohistochemical staining procedures were performed as previously described with minor modifications ^72^.

### Image acquisition and analysis

Fluorescence images were acquired using an Olympus FluoView FV3000 confocal laser scanning microscope equipped with 20× air, 40×, or 63× oil immersion objectives. Imaging parameters were adjusted to ensure detection of immunostained antigen signals above threshold (>150 arbitrary units) while avoiding saturation. All images were acquired under identical exposure settings within each experiment.

For the sciatic nerve (e.g., distal to the crush site), ROIs were defined within the endoneurium, and confocal z-stacks consisting of 3–10 optical sections were acquired and processed as maximum-intensity projections.

For spinal cord analyses, images were consistently collected from the gray matter of the ventral horn. Confocal z-stacks comprising 10–17 optical sections were acquired and collapsed into maximum-intensity projections for analysis. Marker expression and cellular localization were quantified in defined regions of interest (ROIs) within the gray matter of the spinal cord.

Marker expression and cellular localization were quantified in defined regions of interest (ROIs) within the sciatic nerve (e.g., distal to the crush site) and spinal cord using ImageJ software (NIH) and Fiji (an open-source ImageJ distribution).

Fluorescence intensity was quantified by measuring the integrated density (mean gray value) within each ROI and expressed in arbitrary units (a.u.) corresponding to grayscale values of 12-bit images (range: 0–4,095). Quantification also included measurements of positive areas above a defined threshold and/or cell counts as appropriate. The positive area was calculated as the ratio between the area occupied by the marker of interest and the total ROI area.

Quantification was performed blinded to experimental groups where applicable.

### Markers and targets

Immunostaining was performed to assess:

- Axonal regeneration (SCG10, NFL)
- Myeloid/macrophage activation (IBA1, CD68)
- Schwann cells (S100β)
- Myelin markers (MBP; dMBP)
- Lysosomal/autophagy markers (LAMP1, Lysotracker, GPNMB, p62, LC3A/B)

### Human derived iPSC culture

Human iPSCs (WT29 iCas9-Dox TNT434; ^73^ were maintained under feeder-free conditions in mTeSR Plus medium (STEMCELL Technologies) supplemented with 1% penicillin/streptomycin and 200 µg/ml Geneticin on laminin-521 (L521; BioLamina)-coated dishes. Coating was performed in PBS containing Ca²⁺ and Mg²⁺ for 2 h at 37 °C or overnight at 4 °C. Cells were seeded at ∼2 × 10 cells per well (6-well format) and cultured for 3 days prior to downstream applications.

iPSCs were dissociated into single cells with TrypLE (GIBCO) and plated onto laminin-521 coated plates at a seeding density of ∼25000 cells/cm^2^. ROCK inhibitor Y-27632 (10 µM) was added for the first 24 h following seeding, and medium was subsequently changed every second day. Cells were cultured at 37 °C and 5% CO₂.

### CRISPR–Cas9-mediated generation of GPNMB knockout iPSCs

CRISPR–Cas9 genome editing was used to generate GPNMB knockout (KO) iPSC lines. A piggyBAC vector allowing expression of four guide RNAs (gRNAs) targeting distinct regions of the GPNMB coding sequence was stably integrated in the iCas9 iPSCs. The guide sequences were sequence-verified prior to use.

For transfection, 5 × 10 iPSCs were electroporated with 4 µg plasmid DNA and 1 µg transposase using the Lonza Nucleofector 2b system (program B016). Cells were immediately transferred into mTeSR Plus medium supplemented with ROCK inhibitor and seeded onto L521-coated plates. Selection was initiated with puromycin (1 µg/mL) 2 days post-transfection. Selection antibiotics were maintained during culture. Cas9 expression was induced using doxycycline (1 µg/mL) for 7 days.

Edited populations were expanded and validated by sequencing the targeted locus. Independent KO clones were subsequently used for downstream experiments. This approach is based on previously described arrayed CRISPR screening strategies ^74^.

### Embryoid body (EB) formation and iPSCs differentiation into macrophages (iMACs)

iPSC differentiation into macrophages (iMACs) was performed as previously described (Baert et al., 2024). Briefly, iPSCs were dissociated and resuspended in mTeSR Plus medium supplemented with ROCK inhibitor Y-27632 (10 µM) at 6 × 10 cells/mL and seeded into low-adhesion BIOFLOAT 96-well plates (100 µL per well) for embryoid body (EB) formation. On day 2, medium was supplemented with BMP4 (100 ng/mL), stem cell factor (hSCF, 40 ng/mL), and VEGF (100 ng/mL). On day 3, medium was partially replaced with reduced concentrations of BMP4 (50 ng/mL), hSCF (20 ng/mL), and VEGF (50 ng/mL).

By day 5, EBs (400–700 µm diameter) were transferred into macrophage differentiation medium (MDM) supplemented with human M-CSF (100 ng/mL) and IL-3 (25 ng/mL) and seeded into gelatin-coated T75 flasks. From day 8 onwards, culture medium was replaced with X-Vivo 15 medium supplemented with M-CSF and IL-3. Medium was refreshed every 3–4 days, and macrophage-like cells (iMACs) were harvested from the supernatant starting approximately 3-6 weeks after differentiation and for up to 3 months.

### iM0 macrophage differentiation

iMACs collected from the supernatant were further differentiated into unpolarized M0 macrophages (iM0) by culturing in RPMI 1640 medium supplemented with 10% fetal calf serum, sodium pyruvate, HEPES buffer, β-mercaptoethanol, and M-CSF (40 ng/mL). Cells were seeded and cultured for 6–7 days with medium changes every 2–3 days prior to downstream assays.

### Phagocytosis assay

pHrodo™ Red *S. aureus* uptake

Phagocytic activity of iPSC-derived M0 macrophages (iM0) was assessed using pHrodo™ Red *Staphylococcus aureus* Bioparticles™ (Invitrogen, A10010), a pH-sensitive probe that fluoresces upon internalization into acidic compartments (e.g. Lysosomes).

iMAC-derived iM0 macrophages were generated as described above and replated in black 96-well plates at a density of 20 × 10^3^ cells per well. Cells were allowed to adhere and recover prior to treatment.

For treatment conditions, cells were pre-treated with bafilomycin A1 (100 nM) for 2 h to inhibit lysosomal acidification, followed by washout before addition of the phagocytic substrate.

pHrodo™ Red *S. aureus* bioparticles were added at a final concentration of 0.5 mg/mL (10 µL per well). Control conditions included wells containing medium only (background control) and wells containing pHrodo particles without cells.

Phagocytic uptake was monitored over time (up to 48 h) using the IncuCyte® live-cell analysis system (Sartorius), enabling kinetic quantification of fluorescence intensity as a measure of particle internalization and processing. Image acquisition was performed at multiple time points (1 h intervals) under identical settings across conditions, and fluorescence signals were quantified using the IncuCyte analysis software.

The analyzed signals were expressed as total integrated intensity, baseline-corrected, and visualized as the area under the curve (AUC) over a 24 h time to capture uptake kinetics. Experiments were performed using independent biological replicates and included GPNMB knockout (KO) and non-targeting control (NTC) cell lines.

### Myelin isolation and labeling

Myelin debris was isolated from brains of C57BL/6 (B6) mice. Briefly, myelin suspensions were washed twice in ice-cold PBS and collected by centrifugation at 13,000 × g for 10 min.

For fluorescent labeling, myelin was resuspended in 0.1 M sodium bicarbonate buffer (pH 8.4) and adjusted to a concentration of 20 mg/mL. The pH-sensitive amine-reactive dye pHrodo™ iFL Green STP ester (Invitrogen) was prepared as a 10 mM stock solution in anhydrous DMSO and added to the myelin suspension at a final concentration of 100 µM. Samples were incubated for 45 min at room temperature in the dark with gentle mixing.

Following labeling, excess dye was removed by centrifugation (13,000 × g, 10 min) and repeated washes in PBS (three times). The labeled myelin pellet was resuspended in 200 µl sterile PBS and stored at 4 °C in the dark until use.

### Myelin internalization assay

Myelin phagocytosis was assessed using myelin debris isolated from C57BL/6 (B6) mouse brains and labeled with the pH-sensitive amine-reactive dye pHrodo™ iFL Green STP ester (Invitrogen), as described above.

iMAC-derived iM0 macrophages were plated as described and exposed to pHrodo™ Green–labeled myelin (final concentration 1 mg/mL, 25 µL per well). Cells were treated with bafilomycin A1 (100 nM) or the mTOR inhibitor AZD8055 (CCG168, 500 nM), prepared as concentrated stocks in culture medium and added to reach final working concentrations. In selected conditions, treatments were applied prior to or concomitantly with myelin exposure, and cells were incubated for up to 18–24 h.

For live-cell imaging experiments, cells were stained with LysoTracker™ Deep Red (Thermo Fisher Scientific; cat. no. L12492) at 50 nM final concentration for 30 min in the dark, at 37°C, to label acidic compartments and imaged using a Yokogawa CV8000 high-content imaging system. Time-resolved imaging was performed to monitor myelin uptake and intracellular trafficking dynamics.

### Image acquisition and quantitative image analysis

Image acquisition was conducted using automated confocal imaging under identical exposure settings across all conditions, acquiring multiple fields per well. Image analysis was performed using CellPathfinder (Yokogawa). Quantification included total fluorescence intensity of pHrodo™-labeled myelin, colocalization with LysoTracker-positive compartments, and segmentation-based measurements of lysosomal features. Specifically, the total area of myelin signal within segmented lysosomes was quantified, together with the number and total area of acidic LysoTracker-positive lysosomes, and the total fluorescence intensity per cell. These metrics were used as readouts of myelin internalization and lysosomal processing. Experiments were performed using independent biological replicates and included GPNMB knockout (KO) and non-targeting control (NTC) cell lines.

### Immunocytochemistry (ICC)

Human iPSC-derived M0 macrophages (iM0) were seeded onto 96-well imaging plates and allowed to adhere overnight. For lysosomal analysis, cells were treated as indicated and, where applicable, incubated with LysoTracker Deep Red (Thermo Fisher Scientific) at a final concentration of 50 nM for 30 min at 37°C in the dark. LysoTracker fluorescence was measured in-live condition together with labelled myelin used to identify acidic lysosomal compartments and quantify lysosome abundance and cargo– lysosome association.

For lysosomal and autophagy analyses, following treatment, cells were washed with cold phosphate-buffered saline (PBS) and fixed in cold 4% paraformaldehyde (PFA) for 12 min at room temperature. Cells were subsequently washed three times with PBS and permeabilized using Intracellular Staining Permeabilization Wash Buffer (BioLegend; Cat. No. 421002) for 15 min at room temperature. After permeabilization, cells were washed with Cell Staining Buffer (BD Biosciences; Cat. No. 420201) and blocked in PBS containing 3% bovine serum albumin (BSA), 0.1% Tween-20 (PBS-T), and, where indicated, 3% normal serum matched to the host species of the secondary antibody for 60 min at room temperature.

Primary antibodies were diluted in 1% BSA in PBS-T and incubated either for 1 h at room temperature or overnight at 4°C in a humidified chamber. The following primary antibodies were used: rabbit anti-p62/SQSTM1 (Abcam, ab56416, 1:300), rabbit anti-LAMP1 (Abcam, ab208943, 1:100), and goat anti-GPNMB (R&D Systems, AF2550, 1:300). Following incubation, samples were washed three times in PBS for 5 min each.

Cells were then incubated with species-specific fluorophore-conjugated secondary antibodies for 1 h at room temperature in the dark, including donkey anti-rabbit Alexa Fluor 488, goat anti-rabbit Alexa Fluor 546, and donkey anti-goat Alexa Fluor 647 (Invitrogen). Nuclei were counterstained with DAPI (1:1000) for 5–10 min. After final PBS washes, plates were stored in PBS at 4°C protected from light until imaging.

### Image acquisition and quantitative image analysis

Fluorescence images were acquired using either an Olympus FluoView FV3000 confocal microscope (microglia) or a Yokogawa Opera Phenix CV8000 high-content imaging system under identical acquisition settings across experimental conditions. For high-content imaging, two or four fields per well were acquired using automated image capture.

Image analysis was performed using ImageJ/Fiji or CellPathfinder software (Yokogawa). Quantification included measurements of fluorescence intensity (integrated density), positive area above threshold, and cell counts within defined regions of interest. Where applicable, segmentation-based analysis was used to quantify subcellular compartments and marker colocalization. Segmentation-based pipelines were used where appropriate to quantify subcellular compartments and p62–LAMP1 or GPNMB– LAMP1 colocalization. Data were normalized to cell number using automated image segmentation and are presented as fluorescence intensity (integrated density), positive area, puncta counts, or colocalization measurements on a per-cell basis, as indicated in the corresponding figure panels.

### Western blot analysis of autophagy markers

Autophagy signaling and flux were assessed by Western blot analysis of iPSC-derived M0 macrophages (iM0) and microglial cells (iMG), including GPNMB knockout (KO) and non-targeting control (WT) lines. Cells were plated in 6-well plates, expanded for 3 days, and treated for 24 h with autophagy modulators, including vehicle (DMSO), bafilomycin A1 (100 nM), chloroquine (KAE218, 10 µM), mTOR inhibitor AZD8055 (also referred to as CCG168, 500 nM) and Hank’s Balanced Salt Solution (HBSS), to induce nutrient starvation.

Following treatment, cells were lysed in RIPA buffer supplemented with protease and phosphatase inhibitors. Lysates were sonicated, clarified by centrifugation, and protein concentration was determined using a BCA assay. Equal amounts of protein were normalized across samples, and 12.5–15 µg of lysate per sample were resolved by SDS–PAGE on 12% gels for LC3A/B and 4–12% Bis-Tris gradient gels (NuPAGE) for all other proteins, using MOPS running buffer. Proteins were transferred to membranes using the Trans-Blot® Turbo™ Transfer System (Bio-Rad).

Membranes were probed with primary antibodies against autophagy and signaling markers, including LC3A/B (1:1000, Cell Signaling Technology, #12741), p62/SQSTM1 (1:1000–1:2000, Abcam, #ab109012), GPNMB (1:1000, Cell Signaling Technology, #38313), phospho-S6 ribosomal protein (p-S6RP, Ser240/244; 1:1000, Cell Signaling Technology, #5364), total S6RP (1:1000, Cell Signaling Technology, #2217), phospho-4EBP1 (1:1000, Cell Signaling Technology, #2855), total 4EBP1 (1:1000, Cell Signaling Technology, #9644), TFEB and phospho-TFEB (1:1000, Cell Signaling Technology), and LAMP2 (mouse monoclonal, Invitrogen, MA1-205), as indicated. β-actin (1:10,000, Millipore, #A5441) was used as a loading control.

Protein bands corresponding to LC3-I/LC3-II conversion, p62 accumulation, and phosphorylation status of mTORC1 downstream targets (S6RP and 4EBP1), as well as TFEB and lysosomal markers (GPNMB), were used as readouts of autophagic activity and lysosomal function. Band intensities were quantified using standard densitometric analysis and normalized to β-actin.

### Western blot densitometric quantification

Western blot band intensities were quantified using Fiji/ImageJ (National Institutes of Health). For each sample, integrated density values were measured using regions of interest (ROIs) of identical size following local background subtraction. Protein abundance was normalized to β-actin for p62/SQSTM1, LC3, and GPNMB analyses. For LC3 immunoblots, only the lipidated LC3-II species, corresponding to the lower migrating band, was quantified and normalized to β-actin. For p-S6RP and p-TFEB, phosphorylation levels were quantified as the ratio of phosphorylated to total protein (p-S6RP/S6RP and p-TFEB/TFEB, respectively). For 4E-BP1, three phosphorylated bands corresponding to distinct phosphorylation states were quantified individually, and the p-4E-BP1/4E-BP1 ratio was calculated for each band. The mean of the three ratios was used for downstream analyses. Normalized values were expressed relative to the NTC DMSO control condition and are presented as fold change relative to NTC DMSO. Quantified data represent independent biological replicates (n =3, 2 or 1).

### Cathepsin B activity assay (Magic Red)

Lysosomal proteolytic activity was assessed in iPSC-derived M0 macrophages (iM0) using the fluorogenic Magic Red™ Cathepsin B substrate (ImmunoChemistry Technologies), which emits red fluorescence upon cleavage by active cathepsin B in acidic intracellular compartments.

iM0 macrophages were plated in black 96-well plates and exposed to experimental treatments, including lipopolysaccharide (LPS, 100 ng/mL), bafilomycin A1 (BafA1, 100 nM), or the cathepsin B inhibitor CA-074 (20 µM). Where indicated, cells were pre-treated with BafA1 or CA-074 for 1 h prior to substrate addition. In selected conditions, cells were also stimulated with myelin (100 µg/mL) to modulate lysosomal activity.

Magic Red Cathepsin B substrate was reconstituted in DMSO to generate a stock solution and diluted 1:10 in distilled water immediately prior to use. The staining solution was added directly to the culture medium at a final dilution of approximately 1:26 and incubated for 30–60 min at 37 °C protected from light. Cells were optionally counterstained with a nuclear dye (e.g., DAPI or Hoechst) for normalization of cell number.

Live-cell imaging was performed using a Yokogawa CV8000 high-content imaging system, with excitation/emission settings compatible with cresyl violet fluorescence (excitation ∼550–590 nm, emission >610 nm). Time-resolved acquisition was performed to monitor Cathepsin B activity dynamics.

Image analysis was carried out using CellPathfinder (Yokogawa), quantifying mean fluorescence intensity per cell and the area of Magic Red–positive signal above threshold. These measurements were used as readouts of lysosomal enzymatic activity. Fluorescence signals were background-corrected and normalized to control conditions.

Experiments included positive controls for lysosomal activation (LPS treatment) and negative controls (CA-074 inhibition or vehicle-treated cells) to validate assay specificity. All analyses were performed using independent biological replicates and included GPNMB knockout (KO) and non-targeting control (NTC) iM0 macrophages.

### Bulk RNA Sequencing of iPSC-Derived Macrophages

Human induced pluripotent stem cell-derived macrophages (iM0) carrying either a CRISPR-generated GPNMB knockout (GPNMB KO) or the corresponding non-targeting control (NT11) genotype were used for bulk RNA sequencing. Cells were plated at 70,000-100,000 cells per well in 24-well plates and exposed to vehicle control (DMSO), myelin (100 μg/mL), bafilomycin A1 (BafA1), or Hank’s Balanced Salt Solution (HBSS) for 2, 8, or 24 h. Four biological replicates were included for each genotype, treatment, and time point combination. The experimental design was intended to assess the effects of GPNMB deficiency on transcriptional programs associated with lysosomal function, autophagy, phagocytosis, lipid handling, and cellular stress responses.

### RNA Isolation

Total RNA was isolated using a phenol-chloroform extraction followed by column purification. Briefly, cells were lysed directly in QIAzol reagent, and lysates were transferred to 2-mL tubes. Chloroform (200 μL) was added to each sample, followed by vigorous mixing and incubation on ice for 3 min. Samples were centrifuged at 18,000 × g for 15 min to achieve phase separation. The aqueous phase was collected and processed using the miRNeasy Micro Kit (Qiagen) on a QIAcube Connect instrument according to a customized RNA_RNeasyMicro_AnimalTissuesAndCells_LargeSampleDNaseDigest protocol including on-column DNase digestion. RNA was eluted in 18 μL RNase-free water and quantified by spectrophotometry.

### Library Preparation and Sequencing

RNA quality and integrity were assessed using an Agilent Bioanalyzer prior to library preparation, and only samples meeting quality control criteria were included for sequencing. RNA-seq libraries were prepared from 100 ng total RNA using the Illumina Stranded mRNA Preparation Kit (Illumina, #20040534) according to the manufacturer’s instructions (#1000000124518, version 03, June 2022). Libraries were indexed, quality controlled, and sequenced on an Illumina NovaSeq X Plus platform to generate paired-end 50-bp reads, yielding an average of approximately 42 million reads per sample. Base calling was performed using RTA 4.50.12 (NovaSeq X Series Control Software v1.4.0.20115), and FASTQ files were generated using BCL Convert v4.4.6. Sequencing reads were processed using the Exon Quantification Pipeline (EQP, version 2.5)^75^. Reads were aligned to the human reference genome (GRCh38; Ensembl release 107)^76^ using STAR (version 2.7.9a)^77^, and gene-level expression was quantified using the EQP framework.

### Bulk RNA-Seq Data Analysis

Gene expression profiles were generated from bulk RNA sequencing data obtained from GPNMB KO and NT11 macrophages across treatment conditions and time points. Gene-level counts generated by EQP were aggregated to Entrez Gene IDs, retaining the maximum count where multiple Ensembl identifiers mapped to the same Entrez gene. Genes were retained for downstream analyses if expressed at ≥1 count per million (CPM) in at least 75% of samples within each genotype × treatment × time point group. Library sizes were normalized using the trimmed mean of M-values (TMM) method implemented in edgeR.

Quality control was performed using principal component analysis (PCA) of library-size-normalized log2 counts-per-million (log2-CPM) values. Samples deviating by more than three standard deviations from their replicate-group centroid along principal components 1-4 were flagged as candidate outliers but retained for all downstream analyses.

Differential expression analyses were performed using the limma-voom framework with voomWithQualityWeights. A single linear model was fitted incorporating genotype, treatment, and time point effects and their interactions, with RNA integrity number (RIN) included as a continuous covariate. The model was specified as:

∼ Genotype × Treatment × Time point + RIN
using NT11, DMSO, and the 2 h time point as reference levels. Genotype-specific effects within each treatment and time point condition were obtained by linear contrasts followed by empirical Bayes moderation. Genes with a Benjamini-Hochberg adjusted *P* value < 0.05 and an absolute log2 fold change > 0.5 were considered differentially expressed.

Functional enrichment analyses, including Gene Ontology (GO), Gene Set Enrichment Analysis (GSEA), and Gene Set Variation Analysis (GSVA), were used to assess biological processes related to lysosomal biology, autophagy, endolysosomal trafficking, phagocytosis, lipid metabolism, and cellular stress responses.

For unbiased pathway analysis, genes were ranked according to limma moderated *t*-statistics and analyzed using fgsea. Enrichment significance was assessed using Benjamini-Hochberg adjusted *P* values, with adjusted *P* < 0.05 considered significant.

For pathway-focused analyses, GSVA was performed using GO-derived and manually curated lysosomal-and phagocytosis-related gene sets containing at least 10 genes represented in the dataset. Sample-level enrichment scores were calculated from log2-CPM expression values and compared between genotypes within each treatment and time point condition.

To investigate lysosomal regulatory programs, differential expression values were extracted for a manually curated set of genes encoding lysosomal regulators assembled from published literature. Differential expression results were visualized as heatmaps showing log2 fold-change (logFC) values (GPNMB KO versus NT11). Genes were further annotated according to whether they have been reported as rate-limiting regulators of lysosomal function and according to their primary role within lysosomal pathways, including lysosome biogenesis, trafficking, fusion, acidification, cargo degradation, autophagy, receptor-mediated uptake, lysosomal recycling, and lipid metabolism.

Protein-protein interaction (PPI) network analysis was performed using STRING interaction data for the curated lysosomal regulator gene set. Interactions were retrieved using STRINGdb (database version 12.0) with a minimum combined interaction score of 150. Networks were visualized using igraph and ggraph. Nodes represent individual proteins and edges represent known or predicted protein-protein interactions. Node color indicates the direction of differential expression in GPNMB KO macrophages relative to NT11 controls under myelin treatment, and node size is proportional to degree centrality.

### Transcription factor activity inference

Transcription factor (TF) activity was inferred using the decoupleR R package (v2.17.0) and its univariate linear model (ULM) method (run_ulm). This approach regresses a gene-level statistic against the signed mode of regulation for the target genes of each TF and returns a t-value representing inferred TF activity. TF-target interactions were obtained from the human DoRothEA regulon (dorothea_hs), restricted to high-confidence interactions (confidence levels A and B), comprising 118 TFs and 6,408 TF-target interactions.

For the in vitro analysis, the input matrix consisted of genotype-effect moderated t-statistics derived from limma contrasts comparing GPNMB knockout (KO) and non-targeting control (NT11) iPSC-derived macrophages following myelin treatment at 2, 8, and 24 h. No additional statistical model was fitted. The final matrix contained 15,190 genes; when multiple Entrez IDs mapped to the same gene symbol, moderated t-statistics were averaged. TFs represented by fewer than five target genes in the input matrix were excluded (minsize = 5), resulting in 114 TFs retained for analysis. P values were adjusted using the Benjamini-Hochberg method independently for each time point.

Positive activity scores indicate higher inferred TF activity in GPNMB KO cells relative to NT11 controls, whereas negative scores indicate lower inferred activity.

### Cross-validation in an independent in vivo dataset

To validate findings in an independent dataset, the same DoRothEA regulon and ULM framework were applied to differential expression results from the iSNAT sciatic nerve crush dataset. Because moderated t-statistics were not available, genes were ranked using the statistic recommended by decoupleR, defined as −log10(adjusted *p* value) × log2 fold change. Adjusted *p* values were bounded at 1 × 10⁻³ to avoid infinite values. Genes with missing values in any contrast were excluded, yielding a final dataset of 11,502 genes across two post-injury time points (Day 3 and Day 7 relative to Day 0). TF activity was inferred using the same parameters (minsize = 5), and P values were adjusted using the Benjamini-Hochberg procedure within each comparison. TFs with adjusted *p* < 0.01 in the in vitro and in vivo analyses were intersected to identify candidate regulators consistently altered across both datasets.

### Single-Cell RNA Sequencing Analysis of Injured Sciatic Nerve Macrophages

Single-cell RNA sequencing data from mouse sciatic nerve macrophages following peripheral nerve injury (GSE198582) were reanalyzed in R (v4.6.0) using Seurat (v5.5.0) and SeuratObject (v5.4.0). Raw count matrices were normalized using SCTransform (sctransform v0.4.3; glmGamPoi), followed by doublet removal and reclustering, yielding 16,781 high-quality cells. Principal component analysis (PCA) was performed using the first 10 principal components, and cell clustering was conducted using the Leiden algorithm (igraph v2.3.1) at a resolution of 0.2, resulting in eight transcriptionally distinct clusters. Uniform manifold approximation and projection (UMAP) was used for visualization.

Per-cell pathway activity scores were calculated using single-sample Gene Set Enrichment Analysis (ssGSEA) implemented in GSVA (v2.6.5). Gene set collections included Gene Ontology (GO) terms related to autophagy and phagocytosis retrieved from org.Mm.eg.db, as well as curated signatures encompassing lysosome organization, lysosomal lumen, secondary lysosome, autophagosome-lysosome fusion, phagocytosis, phagosome, autophagy, and macroautophagy pathways. Gene sets containing fewer than 10 genes were excluded.

Gene expression patterns were compared across Day 0, Day 3, and Day 7 post-injury samples. Cluster marker genes were identified using Seurat and visualized using ComplexHeatmap (v2.28.0), patchwork (v1.3.2), and ggplot2 (v4.0.3). Disease-associated microglia/macrophage (DAM) signatures were derived from Keren-Shaul et al. (2017) (GSE98971) and evaluated across macrophage clusters using focused gene panels related to lysosomal function, lipid metabolism, antigen presentation, and immune regulation.

### Quantification and statistical analysis

For bulk RNA-seq experiments, differential gene expression analyses were performed using the limma-voom framework with empirical Bayes moderation. A linear model incorporating genotype, treatment, time point, and their interactions, with RIN included as a covariate, was used to estimate genotype-specific effects within each treatment and time point condition. Differential gene expression is reported as log2 fold change (logFC) values of GPNMB KO relative to NT11 samples. Genes with a Benjamini-Hochberg adjusted *P* value < 0.05 and an absolute log2 fold change > 0.5 were considered significantly differentially expressed.

For GSVA analyses, sample-level enrichment scores were calculated from log2-CPM expression values using GO-derived and curated lysosomal- and phagocytosis-related gene sets containing at least 10 represented genes. Enrichment scores were compared between genotypes within each treatment and time point condition using one-way ANOVA followed by Tukey’s honestly significant difference (HSD) post hoc test.

For reanalysis of public transcriptomic datasets, one-way ANOVA followed by Holm-corrected multiple-comparisons tests was used to assess temporal changes in gene expression.

All statistical analyses were performed in R (v4.6.0) and GraphPad Prism 10 (GraphPad Software). Data are presented as mean ± s.d. or mean ± s.e.m., as indicated in the corresponding figure legends. Statistical significance was assessed using one-way or two-way ANOVA, as appropriate, followed by Tukey’s or Sidak’s multiple-comparisons tests. Exact statistical tests, sample sizes, and significance levels are reported in the corresponding figure legends. *P* < 0.05 was considered statistically significant.

---

**Figure S1:**
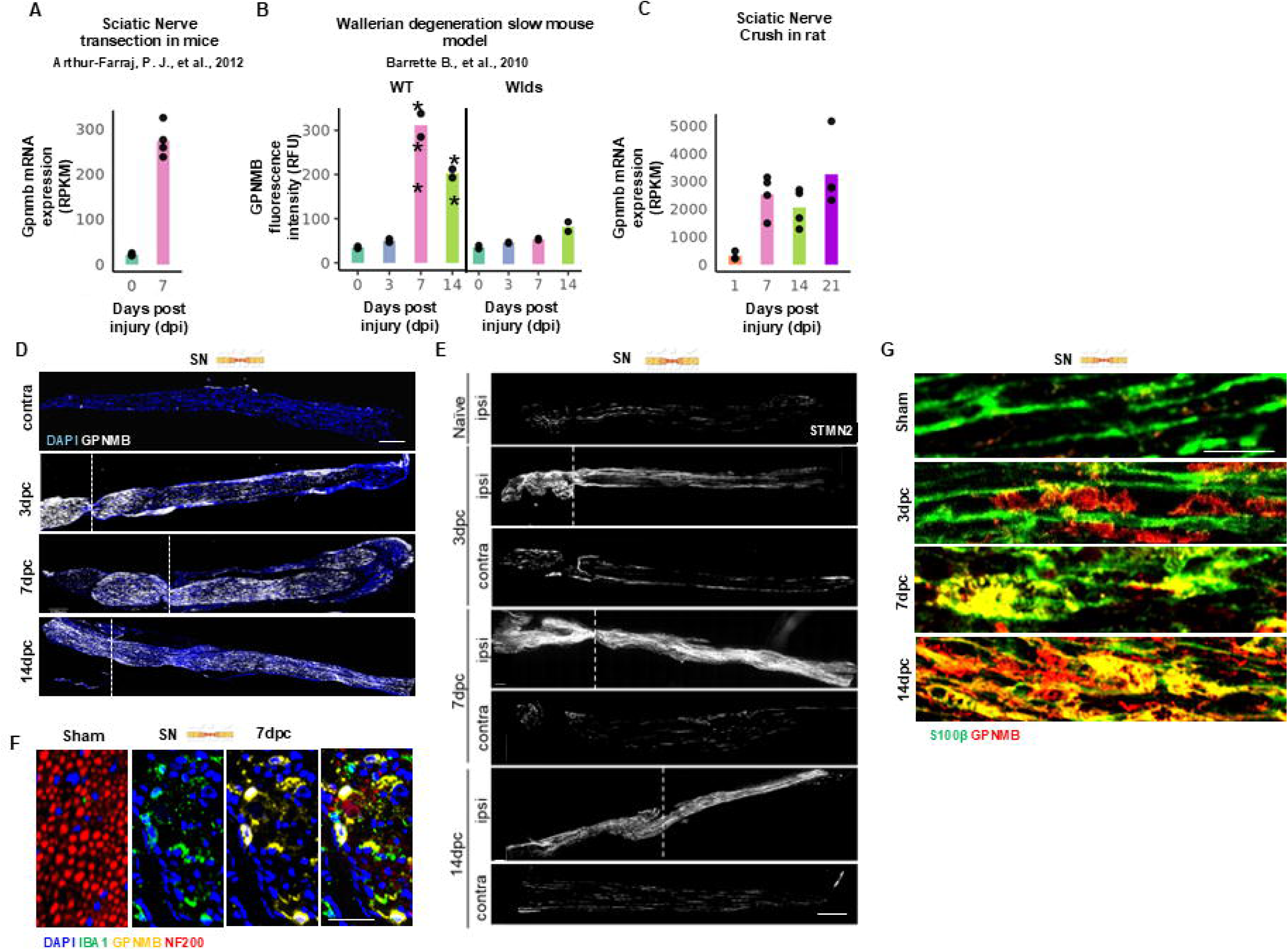
GPNMB is induced after peripheral nerve injury and localizes to macrophages and Schwann cells during regeneration. **(A)** Reanalysis of the peripheral nerve transection dataset reported by Arthur-Farraj et al. (2012) showing *Gpnmb* mRNA expression in uninjured nerves (D0) and at 7 dpc. *Gpnmb* expression was markedly induced following nerve injury. Dots represent individual biological replicates. **(B)** Reanalysis of the mouse sciatic nerve crush microarray dataset from Barrette et al. (2010) showing *Gpnmb* expression in wild-type and Wld^S^ mutant nerves at day 0 and 3, 7, and 14 days post-crush (dpc). Bars represent mean *Gpnmb* microarray signal intensity (RFU), and dots represent individual biological replicates. *Gpnmb* expression increased following injury in wild-type nerves, whereas induction was attenuated in Wld^S^ nerves. **(C)** Reanalysis of an independent rat sciatic nerve injury dataset showing *Gpnmb* mRNA expression during degeneration and regeneration phases (1, 7, 14, and 21 days post-injury). *Gpnmb* expression increased after injury and remained elevated throughout the regenerative response. Dots represent individual biological replicates. **(D)** Representative longitudinal sections of injured sciatic nerves stained for GPNMB (white) and DAPI (blue) at 3, 7, and 14 days post-crush (dpc). Contralateral nerves are shown as uninjured controls. GPNMB expression increased following injury and remained elevated throughout the regenerative phase. Scale bars, 200 µm. **(E)** Representative confocal images of longitudinal 12-μm sciatic nerve sections from C57BL/6 mice stained for STMN2/SCG10 to visualize regenerating axons. Images are shown from naïve nerves, ipsilateral injured nerves (ipsi), and contralateral uninjured nerves (contra). The dashed line indicates the crush site. Scale bars, 200 μm. **(F)** High-magnification confocal images of transverse sciatic nerve sections from sham-operated nerves (left) and injured nerves at 7 days post-crush (dpc) (right), stained for IBA1 (green), GPNMB (yellow), NF200 (red), and DAPI (blue). Arrowheads indicate IBA1⁺ macrophages co-expressing GPNMB. Scale bars, 10 μm. **(G)** Representative high-magnification images of longitudinal sciatic nerve sections from sham-operated (uninjured) nerves and nerves collected at 3, 7, and 14 days post-crush (dpc), stained for GPNMB (red) and the Schwann cell marker S100β (green). Yellow signal in merged images indicates co-localization of GPNMB and S100β. Scale bars, 10 μm. Statistics: Significance indicates pairwise comparisons with the corresponding day 0 control within each genotype using one-way ANOVA followed by Holm-corrected post hoc *t* tests. *p* < 0.05, **p** < 0.01, ***p*** < 0.001.

**Supplementary Figure S2.**
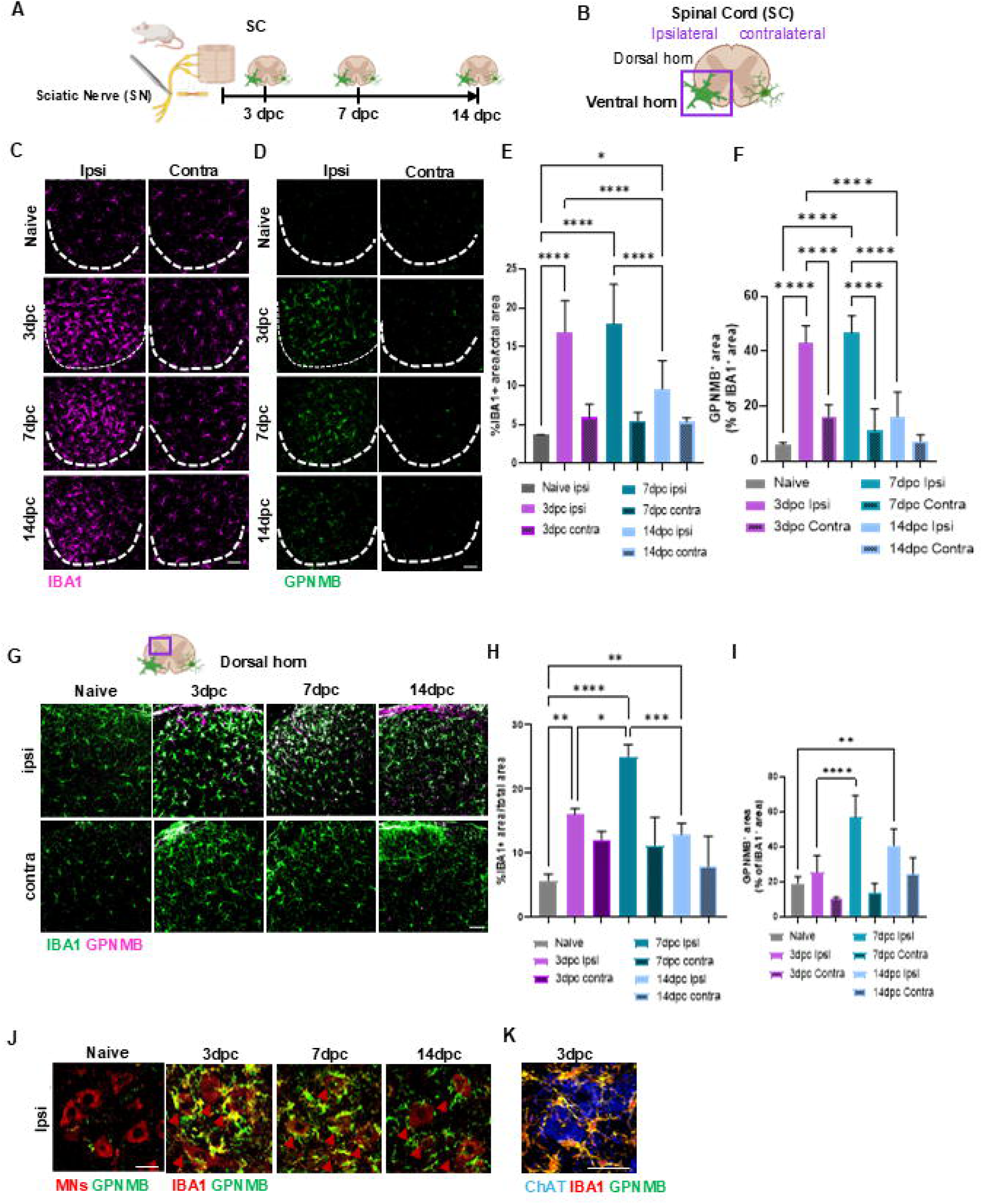
GPNMB is induced in injury-associated myeloid cells in the spinal cord following peripheral nerve injury. **(A)** Experimental design. Schematic of the sciatic nerve crush injury paradigm showing the experimental timeline and collection of lumbar spinal cord tissue at the indicated time points. **(B)** Schematic of the lumbar spinal cord illustrating the ipsilateral and contralateral ventral horn regions analyzed following sciatic nerve crush injury. The boxed region indicates the motor neuron-rich ventral horn area selected for imaging and quantification. **(C–D)** Representative images of lumbar spinal cord ventral horn sections from naïve mice and mice at 3, 7, and 14 days post-crush (dpc) stained for IBA1 (magenta; **C**) or GPNMB (green; **D**). Dashed lines delineate the ventral horn region analyzed. Scale bars, 40 μm. **(E–F)** Quantification of IBA1-positive area **(E)** and GPNMB-positive area **(F)** as a percentage of total ventral horn area in the ipsilateral (injured) and contralateral (uninjured) lumbar spinal cord ventral horns from naïve mice and mice at 3, 7, and 14 days post-crush (dpc). **(G)** Representative confocal images of the dorsal horn region of lumbar spinal cord sections, as indicated by the boxed area in the schematic, from naïve mice and mice at 3, 7, and 14 days post-crush (dpc). Sections were stained for IBA1 (green) and GPNMB (magenta). Scale bars, 40 μm. **(H–I)** Quantification of IBA1-positive area **(H)** and GPNMB-positive area **(I)** as a percentage of total dorsal horn area in ipsilateral and contralateral lumbar spinal cord sections from naïve mice and mice at 3, 7, and 14 days post-crush (dpc). **(J–K)** High-magnification confocal images of the lumbar spinal cord ventral horn following sciatic nerve crush injury. **(J)** Representative images of sham and injured spinal cords at 3, 7, and 14 dpc stained for IBA1 (red) and GPNMB (green). Autofluorescent motoneuron cell bodies (MNs) were used to identify ventral horn motor neurons (red). Arrowheads indicate IBA1⁺GPNMB⁺ microglia clustered around motor neurons, particularly at 3 and 7 dpc. **(K)** Representative confocal image of the ventral horn at 3 dpc stained for ChAT (blue), IBA1 (red), and GPNMB (green), showing GPNMB-expressing microglia surrounding ChAT-positive motor neurons. Scale bars, 20 μm (J) and 10 μm (K). Data are presented as mean ± SD. Statistical significance was assessed using two-way ANOVA followed by Tukey’s multiple comparisons test. Data represent *n* = 5 mice per condition and time point. *\*p < 0.05, **p < 0.01, ***p < 0.001 and ****p < 0.0001*.

**Supplementary Figure S3.**
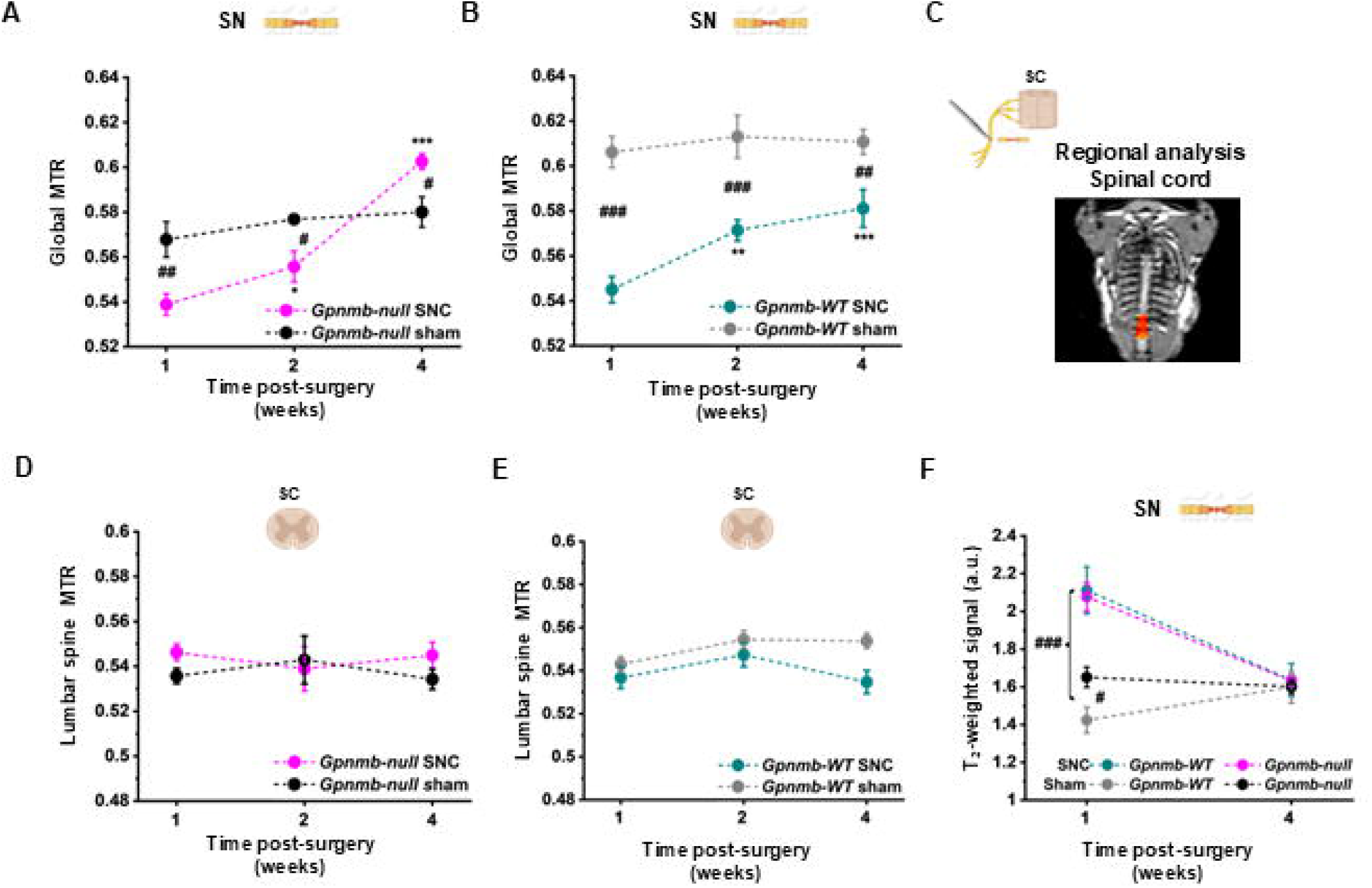
GPNMB deficiency alters MRI measures in the sciatic nerve but not the spinal cord after sciatic nerve crush injury. **(A-B)** Longitudinal assessment of global magnetization transfer ratio (MTR) in the sciatic nerve segment of *Gpnmb*-null (**A**) and *Gpnmb*-WT (**B**) mice following SNC or sham surgery. MTR values are shown for SNC mice and genotype-matched sham-operated controls at 1, 2, and 4 weeks after surgery. **(C)** Schematic illustrating the spinal cord region selected for MRI-based analysis following sciatic nerve crush injury. **(D-E)** Quantification of spinal cord magnetization transfer ratio (MTR) values in sham-operated and SNC mice at the indicated time points after surgery. **(F)** Longitudinal assessment of T2-weighted MRI signal intensity in *Gpnmb*-WT and *Gpnmb*-null mice following sciatic nerve crush (SNC) or sham surgery. Signal intensity was quantified at the indicated time points post-surgery and is shown as mean ± SEM. Statistical analysis was performed using a mixed-effects model with genotype, treatment, time, and their interactions, followed by Fisher’s post hoc test for multiple comparisons. Data are presented as mean ± SEM. Statistical analysis was performed using a mixed-effects model with genotype, time, and their interaction, followed by Fisher’s post hoc test. *P* values are denoted as *\*p < 0.05, **p < 0.01, ***p < 0.001* and *\*\*\*\*p < 0.0001 v*ersus genotype-matched sham controls; *#p < 0.05, ##p < 0.01, ###p < 0.001 and ####p < 0.0001* between genotypes at the indicated time point. Data represent *n* = 10 mice per genotype for SNC and *n* = 5 mice per genotype for sham controls

**Figure S4.**
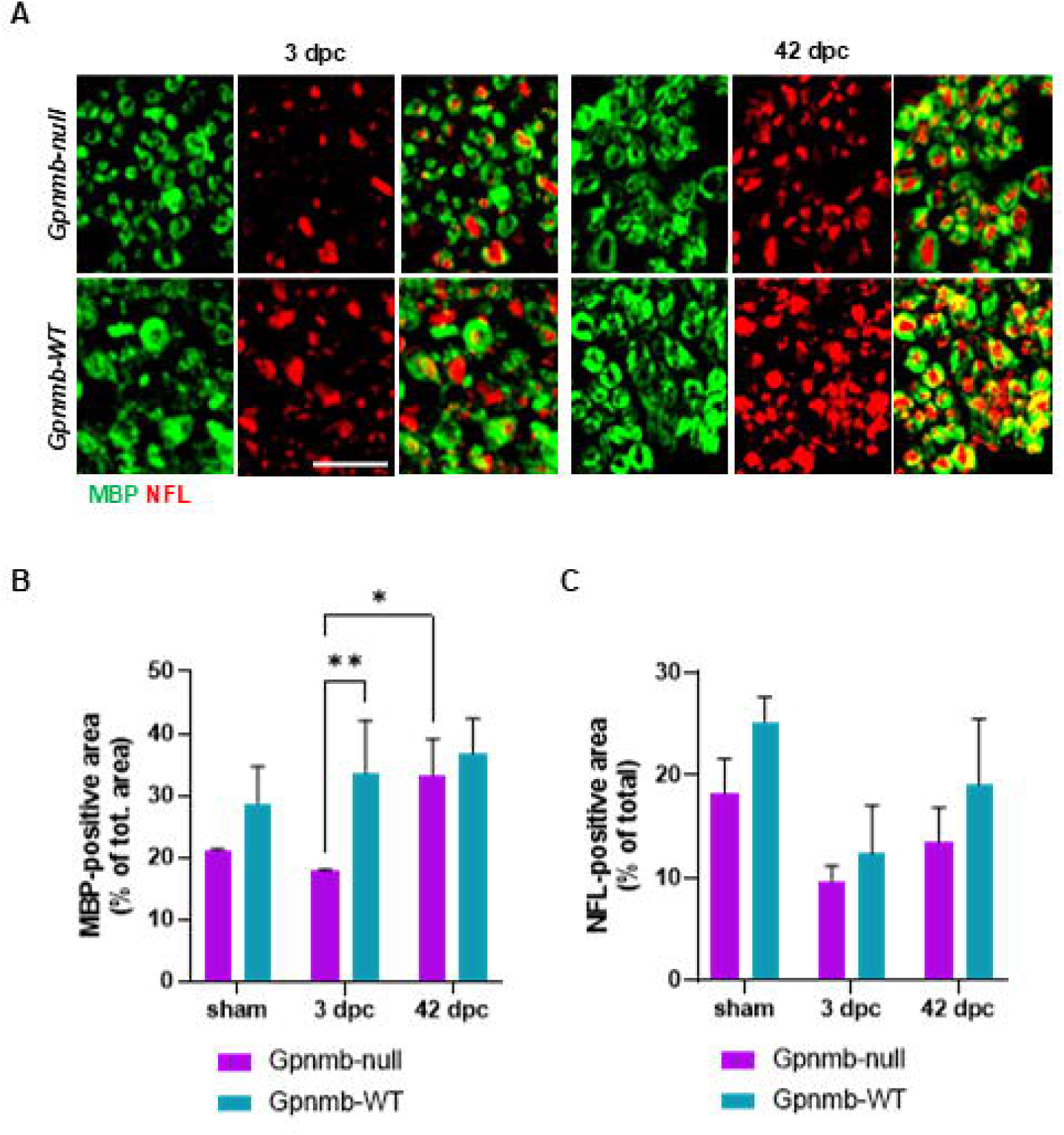
GPNMB promotes myelin debris clearance following peripheral nerve injury. **(A)** Representative high-magnification confocal images of transverse sections of the distal sciatic nerve from *Gpnmb*-null and WT mice at 3 and 42 days post-crush (dpc), stained for myelin basic protein (MBP, green) and neurofilament light chain (NFL, red). Scale bar, 15 μm. **(B-C)** Quantification of MBP-positive area **(B)** and NFL-positive area **(C)** as a percentage of total distal sciatic nerve area in sham-operated mice and in *Gpnmb*-deficient and WT mice at 3 and 42 dpc. Data are presented as mean ± SD. Statistical analyses were performed using a mixed-effects model (REML) with genotype and condition/time (sham, 3 dpc, and 42 dpc) as fixed effects, including the genotype × condition/time interaction, followed by Šídák’s multiple-comparisons test. Multiple images from individual nerves were included in the analysis. Quantification was performed using images acquired from n = 2 mice per genotype in the sham group and n = 3 mice per genotype at each post-injury time point (3 and 42 dpc).

**Figure S5.**
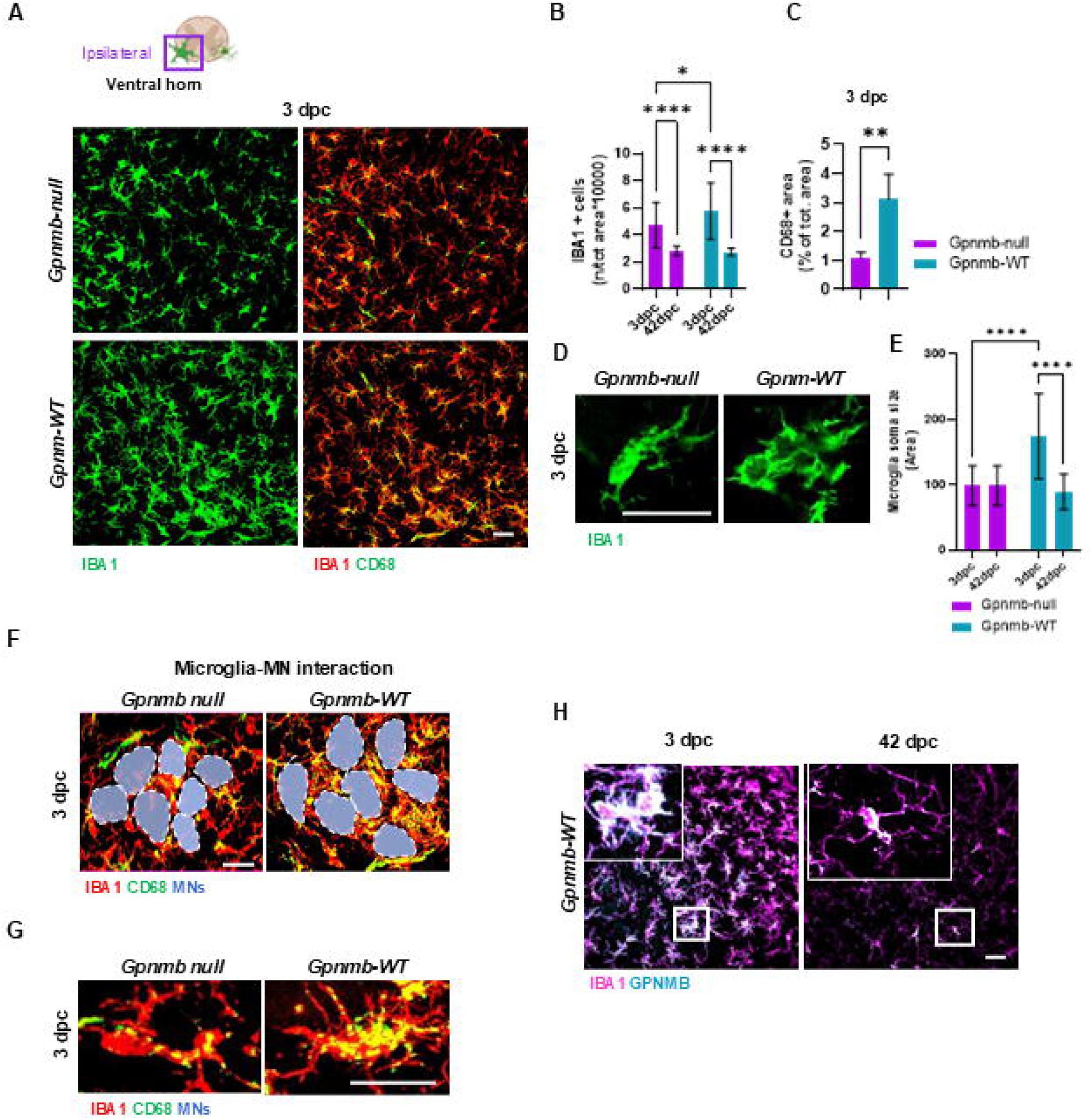
GPNMB regulates microglial activation in the spinal cord following peripheral nerve injury. **(A)** Schematic of the lumbar spinal cord indicating the ipsilateral ventral horn region analyzed following sciatic nerve crush injury (top). Representative confocal images of the ipsilateral ventral horn from *Gpnmb*-null and WT mice stained for IBA1 (green) or co-stained for IBA1 (red) and CD68 (green) (bottom). Scale bar, 40 μm. **(B-C)** Quantification of IBA1-positive microglia at 3 and 42 days post-crush (dpc) (**B**) and CD68-positive area within IBA1-positive microglia at 3 dpc (**C**) in the ipsilateral lumbar spinal cord ventral horn following sciatic nerve crush injury. Data are presented as mean ± SD. For (**B)**, statistical analysis was performed using two-way ANOVA followed by Tukey’s multiple comparisons test. For **(C)**, statistical significance was assessed using Welch’s *t* test. **(D)** Representative high-magnification confocal images of IBA1-positive microglia in the ipsilateral ventral horn from *Gpnmb*-null mice and WT 3 dpc. Images highlight injury-associated morphological changes in microglia and were used for quantitative morphological analysis. Images represent maximum-intensity projections generated from three optical sections acquired at 1-μm intervals. Scale bar, 5 μm. **(E)** Quantification of microglial soma area in the ipsilateral ventral horn 3 dpc. Soma size was measured from IBA1-positive microglia. Data are presented as mean ± SD. Statistical significance was assessed using two-way ANOVA followed by Tukey’s multiple comparisons test. **(F)** Representative high-magnification confocal images of the ipsilateral ventral horn 3 dpc, showing IBA1-positive microglia (red) with CD68-positive lysosomal compartments (green) closely associated with motor neurons (gray outlines). Images illustrate the accumulation of CD68-enriched microglia around motor neuron somata in the injured ventral horn. Scale bar, 15 μm. **(G)** Representative high-magnification confocal images of individual IBA1-positive microglia in the ipsilateral ventral horn from *Gpnmb*-null and WT mice 3 dpc. CD68-positive lysosomal compartments are shown in green and IBA1-positive microglial processes in red. Images highlight the intracellular distribution of CD68-positive lysosomes within activated microglia. Scale bar, 5 μm. **(H)** Representative confocal images of the ipsilateral ventral horn at 3 and 42 dpc stained for IBA1 (magenta) and GPNMB (cyan). Higher-magnification insets of the boxed regions illustrate the dynamic changes in GPNMB expression within IBA1-positive microglia over the course of the injury response. Scale bars, 40 μm (main images) and 20 μm (insets). Data represent *n* = 5 mice per genotype. Statistical significance is denoted as \**P* < 0.05, \*\**P* < 0.01, \*\*\**P* < 0.001, and \*\*\*\**P* < 0.0001.

**Figure S6.**
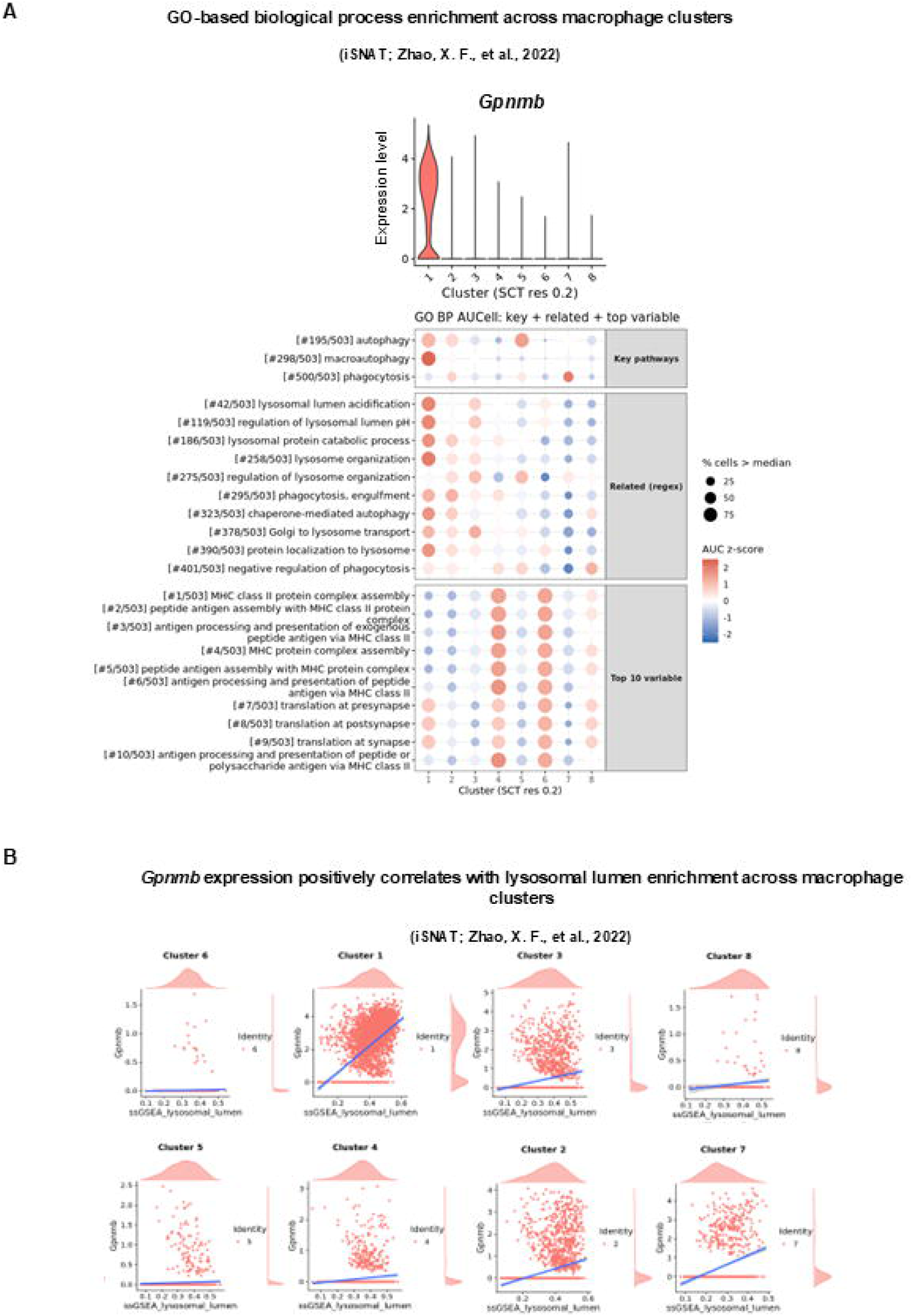
GPNMB marks injury-associated macrophages enriched for phagocytic, lysosomal, and lipid-processing programs during sciatic nerve regeneration. **(A)** Top, violin plot showing *Gpnmb* expression across macrophage clusters identified by single-cell RNA sequencing of injured sciatic nerves from the previously published injured sciatic nerve atlas (iSNAT) dataset (Zhao et al., 2022). Bottom, AUCell-based pathway enrichment analysis of macrophage clusters from the same dataset. Dot size indicates the percentage of cells with AUCell scores above the median, and color indicates the scaled AUCell score (z-score). The *Gpnmb*-expressing macrophage cluster is enriched for gene programs associated with phagocytosis, lysosomal function, autophagy, and lipid processing. **(B)** Scatter plots showing *Gpnmb* expression as a function of ssGSEA lysosomal lumen enrichment scores across macrophage clusters identified by single-cell RNA sequencing. Cluster 1, the *Gpnmb*-high macrophage cluster, exhibits a positive association between *Gpnmb* expression and lysosomal lumen enrichment. Blue lines indicate linear regression fits. Marginal density plots show the distribution of values for each variable.

**Figure S7.**
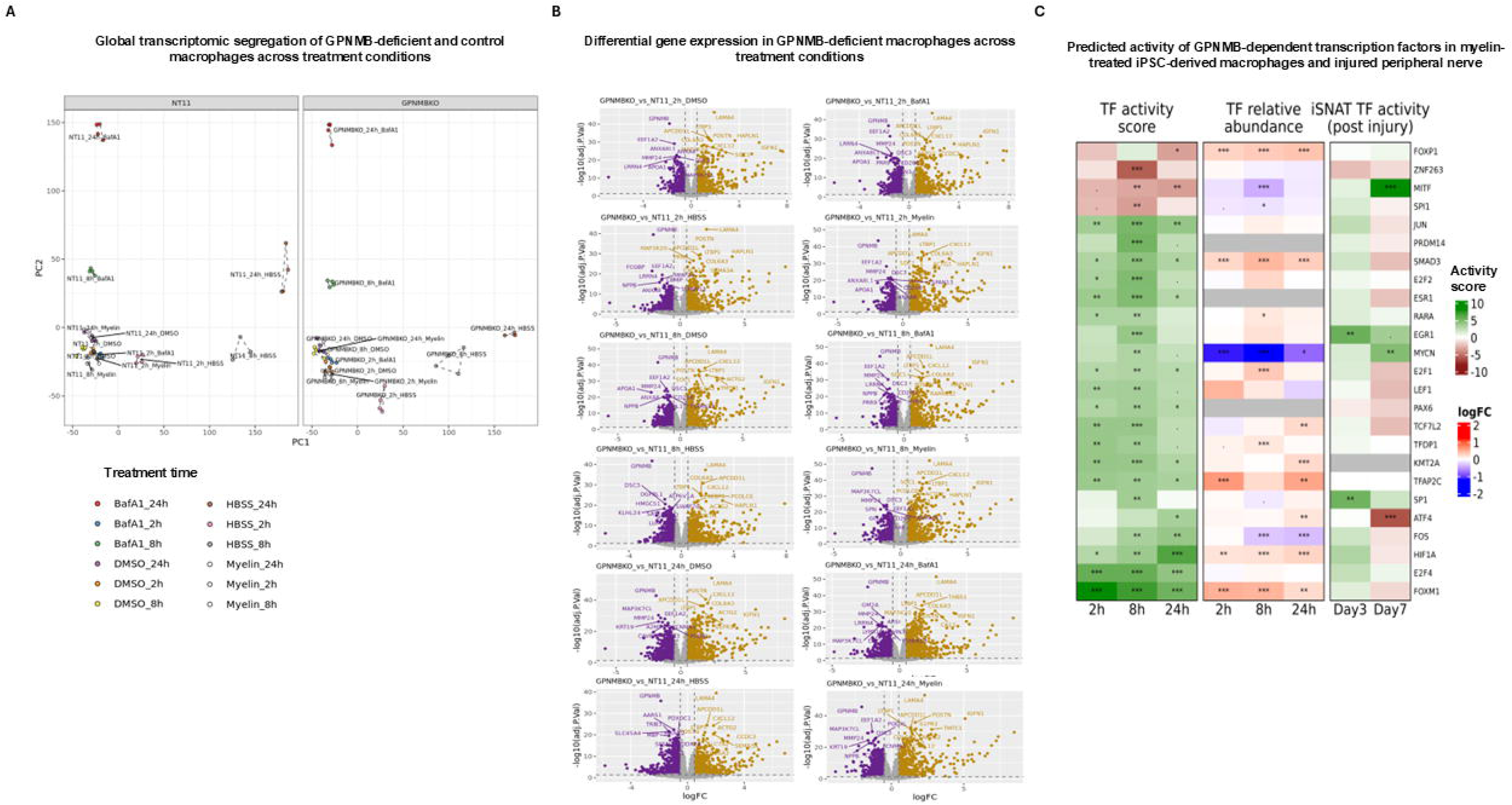
Bulk RNA-seq reveals transcriptional and regulatory programs altered by GPNMB deficiency in iPSC-derived macrophages. **(A)** Principal component analysis (PCA) of bulk RNA-seq samples from non-targeting control (NT11) and GPNMB knockout (KO) iPSC-derived macrophages treated with DMSO, HBSS, Bafilomycin A1 (BafA1), or myelin for 2, 8, or 24 h. Samples cluster according to treatment and genotype. **(B)** Volcano plots showing differential gene expression between GPNMB KO and NT11 control macrophages under each treatment condition and time point. Genes meeting differential expression significance thresholds are highlighted. Due to plot density, only a subset of significantly differentially expressed genes that could be clearly displayed were annotated. **(C)** Predicted transcription factor (TF) activity in myelin-treated GPNMB knockout (KO) versus NT11 control iPSC-derived macrophages and in post-injury macrophages from the iSNAT dataset. Rows represent the 25 TFs showing the strongest genotype-dependent activity changes in vitro (smallest Benjamini-Hochberg [BH]-adjusted *p* value across myelin time points), ordered by hierarchical clustering of in vitro TF activity scores and displayed in the same order across all panels. Left, inferred TF activity in GPNMB KO relative to NT11 macrophages following myelin treatment for 2, 8, and 24 h (green, higher activity in GPNMB KO; red, lower activity in GPNMB KO). Middle, differential expression of the corresponding TF transcripts in the same comparison (red, upregulated in GPNMB KO; blue, downregulated in GPNMB KO). Right, inferred TF activity in macrophages from the iSNAT sciatic nerve crush dataset at day 3 and day 7 post injury relative to day 0, shown on the same activity scale. Color scales are centered on zero and scaled independently for each panel. Blank cells indicate TFs absent from the corresponding dataset. Asterisks indicate BH-adjusted *p* values (\*\**p* < 0.001, \**p* < 0.01, *p* < 0.05, · *p* < 0.1). See Materials and Methods for details of TF activity inference.

**Figure S8.**
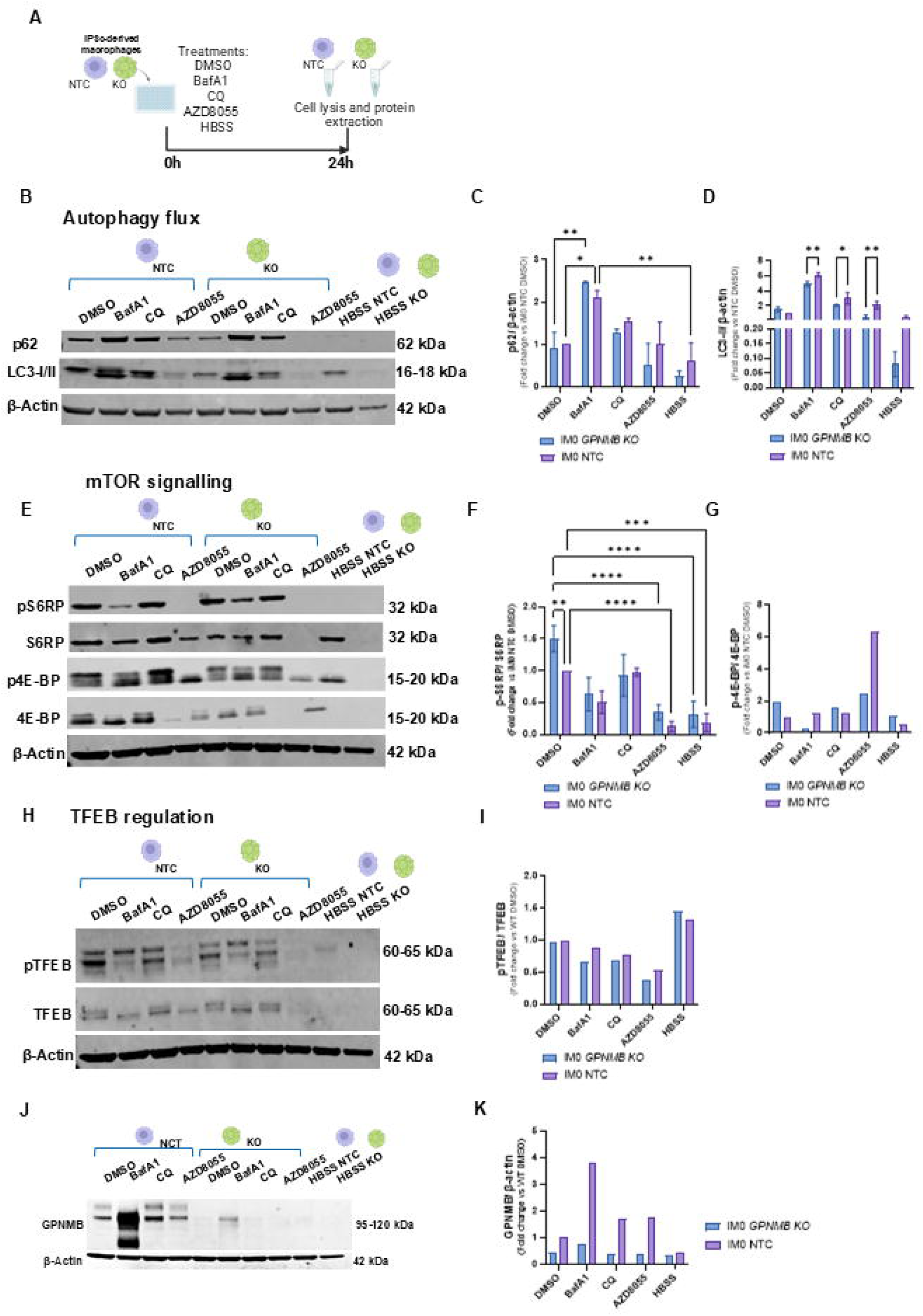
GPNMB supports lysosomal homeostasis and autophagic function in human macrophages. **(A)** Experimental design. Human iPSC-derived macrophages (NTC and *GPNMB* KO) were treated for 24 h with DMSO, bafilomycin A1 (BafA1; lysosomal inhibitor), chloroquine (CQ; lysosomal inhibitor), AZD8055 (ATP-competitive mTOR kinase inhibitor targeting both mTORC1 and mTORC2), or HBSS starvation medium, followed by protein extraction for immunoblot analysis. **(B)** Immunoblot analysis of autophagy flux in human iPSC-derived NTC and *GPNMB* KO macrophages following 24 h treatment with DMSO, bafilomycin A1 (BafA1), chloroquine (CQ), AZD8055, or HBSS. Membranes were probed for p62 and LC3-I/LC3-II. β-actin served as a loading control. **(C-D)** Quantification of p62/SQSTM1 (**B**) and LC3-II (**C**) normalized to β-actin in GPNMB KO and NTC iPSC-derived macrophages following 24 h treatment with DMSO, BafA1, AZD8055, or HBSS. Data are expressed as fold change relative to NTC DMSO. **(E)** Immunoblot analysis of mTOR signaling in NTC and *GPNMB* KO macrophages following the indicated treatments. Membranes were probed for phosphorylated and total S6RP and 4EBP. β-actin served as a loading control. **(F-G)** Quantification of p-S6RP normalized to total S6RP (**F**) and p-4E-BP1 normalized to total 4E-BP1 (**G**) in GPNMB KO and NTC iPSC-derived macrophages following 24 h treatment with DMSO, BafA1, AZD8055, or HBSS. Data are expressed as fold change relative to NTC DMSO. (H) Immunoblot analysis of TFEB regulation in NTC and *GPNMB* KO macrophages following the indicated treatments. Membranes were probed for phosphorylated TFEB (pTFEB) and total TFEB. β-actin served as a loading control. (I) Quantification of p-TFEB normalized to total TFEB in GPNMB KO and NTC iPSC-derived macrophages following 24 h treatment with DMSO, BafA1, AZD8055, or HBSS. Data are expressed as fold change relative to NTC DMSO. **(J-K)** Representative immunoblots of GPNMB and β-actin in GPNMB KO and NTC human iPSC-derived macrophages following 24 h treatment with DMSO, BafA1, CQ, AZD8055, or HBSS. β-actin served as a loading control (**J**). (**K**) Quantification of GPNMB protein abundance normalized to β-actin in GPNMB KO and NTC human iPSC-derived macrophages following 24 h treatment with DMSO, BafA1, CQ, AZD8055, or HBSS. Values are shown relative to DMSO-treated NTC cells. A residual immunoreactive band was detected in GPNMB KO cells, likely reflecting recognition of a truncated product generated by the Aguzzi targeting construct. Data are presented as mean ± SD. Statistical analyses were performed using two-way ANOVA followed by Tukey’s multiple comparisons test unless otherwise indicated. *p* < 0.05, \**p* < 0.01, \*\**p* < 0.001, \*\*\**p* < 0.0001, \*\*\*\**p* < 0.0001.

**Figure S9.**
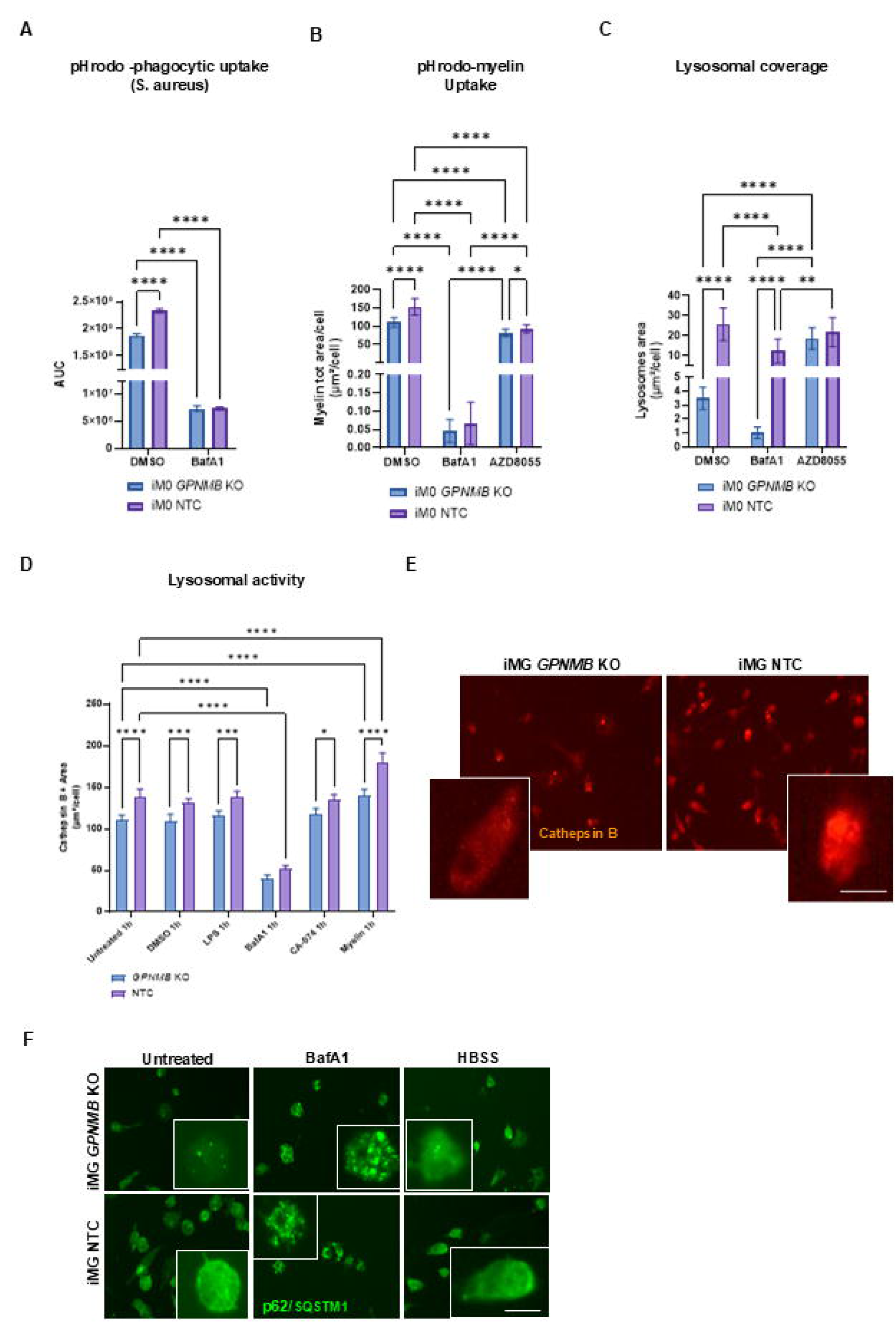
GPNMB deficiency impairs lysosomal function and autophagosome– lysosome fusion during myelin challenge. **(A)** Phagocytic uptake of pHrodo Red-labeled S. aureus bioparticles by human iPSC-derived M0 macrophages (iM0) treated with DMSO or bafilomycin A1 (BafA1). Phagocytic activity is presented as area under the curve (AUC) of fluorescence intensity values normalized to baseline (t = 0). Data are presented as mean ± SD. Statistical analysis was performed using two-way ANOVA followed by Tukey’s multiple-comparisons test. **(B–C)** Quantification of myelin processing in GPNMB KO and NTC iM0 macrophages following uptake of pHrodo-labeled myelin. **(B)** Internalized myelin quantified as total myelin-positive area per cell. **(C)** Lysosomal compartment size quantified as LysoTracker-positive area per cell. Data are presented as mean ± SD. Statistical significance was assessed using two-way ANOVA followed by Tukey’s multiple comparisons test. **(D)** Quantification of lysosomal proteolytic activity in GPNMB KO and NTC iM0 macrophages under untreated conditions or following treatment with DMSO, LPS, bafilomycin A1 (BafA1), CA-074, or myelin. Cathepsin B activity was measured using the Magic Red Cathepsin B reporter assay and quantified as Magic Red-positive area per cell 1 h after probe addition. BafA1 and CA-074 were included as controls for inhibition of lysosomal acidification and Cathepsin B activity, respectively. Data are presented as mean ± SD. Statistical significance was assessed using two-way ANOVA followed by Tukey’s multiple comparisons test. **(E)** Representative fluorescence images of NTC and GPNMB KO iM0 macrophages stained for Cathepsin B activity using the Magic Red reporter assay. Insets show higher-magnification views of individual cells. Scale bars, 10 μm. **(F)** Representative fluorescence images of NTC and GPNMB KO iPSC-derived microglia stained for p62/SQSTM1. Insets show higher-magnification views of individual cells illustrating the distribution and morphology of p62/SQSTM1-positive puncta. Scale bars, 20 μm (main images) and 10 μm (insets). Data are presented as mean ± SD. Statistical significance was assessed using two-way ANOVA followed by Tukey’s multiple comparisons test. Data represent *n* = 2-5 independent experimental replicates. Statistical significance is denoted as \**P* < 0.05, \*\**P* < 0.01, \*\*\**P* < 0.001, and \*\*\*\**P* < 0.0001.

